# A microbial derived bile acid acts as GPBAR1 agonist and RORγt inverse agonist and reverses inflammation in inflammatory bowel disease

**DOI:** 10.1101/2024.04.08.588556

**Authors:** Michele Biagioli, Cristina Di Giorgio, Carmen Massa, Silvia Marchianò, Rachele Bellini, Martina Bordoni, Ginevra Urbani, Rosalinda Roselli, Ginevra Lachi, Elva Morretta, Fabrizio Dal Piaz, Bruno Charlier, Bianca Fiorillo, Bruno Catalanotti, Luigi Cari, Giuseppe Nocentini, Patrizia Ricci, Eleonora Distrutti, Valentina Sepe, Angela Zampella, Maria Chiara Monti, Stefano Fiorucci

## Abstract

The interplay between the dysbiotic microbiota and bile acids is a critical determinant for development of a dysregulated immune system in inflammatory bowel disease (IBD). Here we have investigated the fecal bile acid metabolome, gut microbiota composition, and immune responses in IBD patients and murine models of colitis and found that IBD associates with an elevated excretion of primary bile acids while secondary, allo- and oxo- bile acids were reduced in both human and mice models of IBD. These changes correlated with the disease severity, mucosal expression of pro-inflammatory cytokines and chemokines, and reduced inflow of anti-inflammatory macrophages and Treg in the gut. Analysis of bile acids metabolome in the feces allowed the identification of five bile acids: 3-oxo-DCA, 3-oxo-LCA, allo-LCA, iso-allo-LCA and 3-oxo-UDCA, whose excretion was selectively decreased in IBD patients and diseased mice. By transactivation assay and docking calculations all five bile acids were shown to act as GPBAR1 agonists and RORγt inverse agonists, skewing Th17/Treg ratio and macrophage polarization toward an M2 phenotype. In a murine model of colitis, administration of 3-oxo-DCA suffices to reverse colitis development and intestinal dysbiosis in a GPBAR1-dependent manner. *In vivo* administration of 3-oxo-DCA to colitic mice also reserves disease severity and RORγt activation induced by a RORγt agonist and IL-23, a Th17 inducing cytokine. These results demonstrated intestinal excretion of 3-oxoDCA, a dual GPBAR1 agonist and RORγt inverse agonist, is reduced in IBD and models of colitis and its restitution protects against colitis development, highlighting a potential role for this agent in IBD management.

## Introduction

Inflammatory bowel disease (IBD) encompassing Crohn’s disease (CD) and ulcerative colitis (UC)^1^ are chronic relapsing inflammatory disorders of the intestine whose incidence and prevalence has increased significantly globally in the last two decades^1^.

While it is generally appreciated that, in genetically predisposed individuals, a dysfunctional regulation of both innate and adaptive immunity plays a role in the development of the abnormal inflammatory response, mechanisms that promotes immune activations are not completely elucidated^2–5^. The innate immune system comprises structural barriers, receptors capable of recognizing conserved motifs in microorganisms, and various cell types, including neutrophils, dendritic cells (DCs), monocytes, macrophages, and natural killer (NK) cells. Together, these components promote a rapid and effective inflammatory response against microbial determinants^6^. This response not only serves as the initial line of defense against infections but also contributes to the activation of T cells through cytokines production and antigens presentation. Thus, while development of CD is mainly associated with aTh1 response, UC is more closely associated with a non-conventional Th2 response^7^. In addition, both CD and UC witnesses a robust involvement of Th17 cells, a subset of inflammatory T cells that proliferate in response to myeloid-derived interleukin (IL)-23^8^.

In homeostatic conditions, the intestinal microbiota plays an important role in modulating the intestinal immune response and several microbiota-derived metabolites, i.e. short-chain fatty acids (SCFAs) (butyric acid, propionic acid, and acetate) and secondary bile acids, promote the development of a tolerogenic phenotype of the intestinal immune system toward microbial determinants^9–11^. Bile acids are atypical steroids generated in the liver from the conversion of cholesterol into two primary bile acids, cholic acid (CA) and chenodeoxycholic acid (CDCA), that are secreted in the biliary tract after conjugation with glycine and taurine and then released into the intestine. In the small intestine, CA and CDCA become the substrates for various bacterial enzymes, allowing their bio-transformations to secondary bile acids, lithocholic acid (LCA), deoxycholic acid (DCA)^12^, hyocholic acids(HCAs) and 3-, 7-, and 12 oxo-bile acid analogues that represent a significant proportion of gut microbiota derived bile acids in the colon^12,13^.

In addition to their function in nutrients absorption, bile acids function as endogenous ligands for membrane and nuclear receptors^14–16^. These receptors, are essential for maintaining lipid and bile acids homeostasis, intestinal barrier integrity and a tolerogenic phenotype of intestinal immune cells^17^ ^15^. GPBAR1, also known as TGR5 (Takeda G protein-coupled receptor 5), is a G protein-coupled receptor (GPCR) that functions as a membrane receptor for DCA and LCA. In the intestine, GPBAR1 is expressed by intestinal epithelial and endocrine L cells, neurons, smooth muscle cells ^14^. GPBAR1 is also expressed by cells of innate immunity and plays a role in modulating the innate immune response and barrier integrity as demonstrated by the fact that Gpbar1^-/-^ mice spontaneously develop an intestinal inflammation with age and are more prone to develop a severe disease in models of IBD^18^. Additionally, expression of GPBAR1 is dysregulated in patients and animal models of UC, but precise mechanisms that regulate GPBAR1 function in IBD have been not elucidated.

The 3-, 7- and 12-oxo-bile acids, generated by the intestinal microbiota, have recently attracted attention because their potential to function as Retinoic Acid Receptor-Related Orphan Receptor (ROR)γt inverse agonists. RORγt is highly represented in Th17 and innate lymphoid cells (ILC)-3 and represent a well-defined therapeutic target in IBD ^1,15,19–23^. In T cells, RORγt agonism promotes the differentiation from naive T cells toward Th17 cells whose activation promotes IBD development in human^24^.

Since, decoding the bile acids/immune cells communications might reveal novel therapeutic approaches in IBD, we have embarked in a project designed to identify whether microbiota-derived bile acids function as endogenous modulators of the intestinal immunity and whether a dysregulated production of these mediators might contribute to development of IBD.

## Results

### Correlation between bile acids, intestinal microbiota, and severity of IBD

We have first investigated whether development of CD and UC associated with an abnormal bile acid pattern in the feces. For this purpose, we have employed the IBD cohort from the Human Microbiome Project (HMP2). The results obtained demonstrated that both CD and UC patients exhibited an altered primary bile acid and its derivatives pattern (Fig. 1a, c), while the secondary bile acids were not modulated in a statistically dependent manner. (Fig. 1b, d). However, in comparison to the healthy controls, both CD and UC displayed a robust increase in the ratio of primary/secondary bile acids (Fig. 1e). Furthermore, the analysis of oxo- and allo- derivatives showed a trend towards a decrease in 3-oxo-DCA and 3-oxo-LCA (Fig. 1f).

**Fig. 1.**
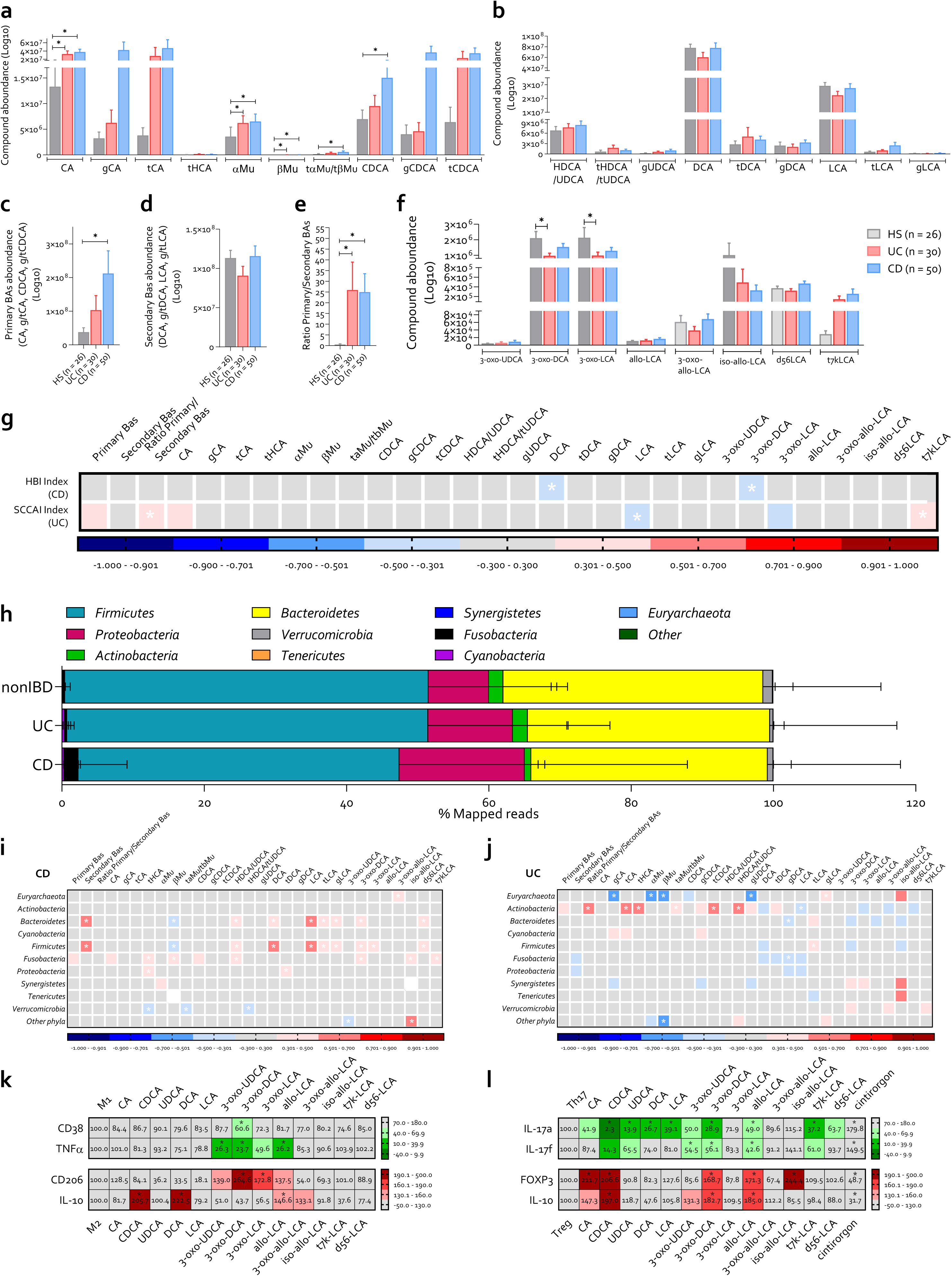
Fecal bile acid alterations and intestinal dysbiosis in CD and UC. We utilized the Inflammatory Bowel Disease (IBD) cohort from the Human Microbiome Project (HMP2) to investigate fecal bile acids and intestinal microbiota in healthy individuals, patients with ulcerative colitis (UC), and patients with Crohn’s disease (CD). Fecal bile acid concentrations: (**a**) primary and conjugated bile acids, (**b**) secondary and conjugated bile acids, (**c**) sum of all primary bile acids, (**d**) sum of all secondary bile acids, (**e**) ratio of primary to secondary bile acids, and (**f**) oxo- and allo-derivatives of bile acids. (**g**) Correlation matrix between fecal bile acid concentrations and the clinical disease index measured using the HBI for CD and the SCCAI for UC. (**h-j**) Analysis of fecal microbiota composition: (**h**) relative abundance of phyla and correlation matrix between fecal bile acid concentration and phyla percentage in (**i**) CD patients and in (**j**) UC patients. Evaluation of the activity of bile acids (1 µM) on the macrophages and T cells polarization: relative mRNA expression of (**k**) CD38 and TNFα in pro-inflammatory M1 macrophages, and CD206 and IL-10 in anti-inflammatory M2 macrophages; (**l**) relative mRNA expression of IL-17a and IL-17f in Th17 cells, and FOXP3 and IL-10 in Treg cells. qPCR data are normalized to GAPDH mRNA and data shown are the mean of 5 replicates for each experimental condition. Abbreviations: BAs, bile acids; HS, healthy samples; UC, ulcerative colitis; CD, Crohn’s disease; HBI, Harvey-Bradshaw Index; SCCAI, Simple Clinical Colitis Activity Index; T, tauro; LCA, lithocholic acid; DCA, deoxycholic acid; CDCA, chenodeoxycholic acid; HDCA, hyodeoxycholic acid; UDCA, ursodeoxycholic acid; HCA, hyocholic acid; αMu, alpha-muricholic acid; βMu, beta-muricholic acid; 7k, 7keto; CA, cholic acid; Dex, dexamethasone; CD38, cluster of differentiation 38; TNFα, tumor necrosis factor-alpha; CD206, cluster of differentiation 206; IL-10, interleukin-10; IL-17a, interleukin-17a; IL-17f, interleukin-17f; FOXP3, Forkhead Box P3.

A correlation study between bile acids concentrations and disease severity, measured using the Harvey-Bradshaw Index (HBI) for CD and the Simple Clinical Colitis Activity Index (SCCAI) for UC, revealed few correlations (Fig. 1g). Nevertheless, CD severity was inversely correlated with DCA and 3-oxo-DCA fecal concentrations. In contrast, UC severity was directly correlated with the total concentration of t7kLCA and the primary/secondary bile acids ratio, and inversely related with LCA. The analysis of gene expression for various cytokines and chemokines in the rectum of these patients showed an up-regulation of these genes compared to the levels measured in non IBD patients and a positive correlation of these biomarkers with disease severity indexes (fig. S1a-d). Furthermore, as shown in fig. S1d, fecal content of secondary bile acids inversely correlated with the expression of various cytokines and chemokines in UC, while there was a direct correlation between IFNγ gene expression and some chemokines with the primary/secondary bile acids ratio. Additionally, dissecting the immune pattern by immune deconvolution of RNAseq data, we demonstrated that disease severity of both CD and UC inversely correlated with the development of M2 anti-inflammatory macrophages (fig. S1e-k).

We have then attempted to correlate the composition of intestinal microbiota with bile acid patterns in IBD patients from the same cohort (Fig. 1h-j). The results of these investigations demonstrated that, while there is a robust variation of the intestinal microbiota structure among patients subsets^25–27^, UC patients were characterized by a slight expansion of *Proteobacteria* and *Actinobacteria* phyla, with a decrease in *Tenericutes* and *Verrucomicrobia*(Fig. 1h). In contrast, CD patients exhibited a more severe dysbiosis characterized by a decrease in *Firmicutes, Bacteroidetes, Tenericute* and *Verrucomicrobia*, along with an increase in *Proteobacteria and Fusobacteria* (Fig. 1h). These alterations in the composition of the intestinal microbiota correlated with changes of fecal bile acids (Fig. 1i, j). Thus, in CD patients, a decrease in *Bacteroidetes* and *Firmicute* positively correlated with decreased excreti0n of DCA, LCA and their oxo- and allo- derivatives (Fig. 1i), while in UC patients, increased abundance of *Actinobacteria* positively correlated with a rise excretion of primary bile acids and inversely correlated with a decrease of secondary bile acids and their oxo- and allo- derivatives (Fig. 1j). Collectively, these data demonstrated that a distinctive feature of IBD is the reduced formation of secondary bile acids and their oxo- and allo- derivatives.

To investigate whether bile acid alterations could promote an immune dysfunction *per se*, we have isolated macrophages and T cells from buffy coats of healthy donors and polarized them toward a M1 or M2 macrophages (Fig. 1k) and Th17 or Treg cells (Fig. 1 l) in the presence or absence of various bile acids. The results of these experiments demonstrated that the 3-oxo-DCA robustly down-regulated the expression of the CD38 and TNFα on M1 macrophages, while the allo-LCA, 3-oxo-LCA and 3-oxo-UDCA down-regulated only the expression of TNFα (Fig. 1k upper panel). Furthermore, exposure of M2 macrophages to allo-LCA, 3-oxo-DCA, 3-oxo-LCA, and 3-oxo-UDCA further increased the expression of the CD206. On the other hand, the expression of IL-10 was positively modulated only by DCA, CDCA, allo-LCA and 3-oxo-allo-LCA (Fig. 1k lower panel).

In contrast, several bile acids effectively reduced the expression of IL-17a in Th17 cells, while only CDCA, UDCA, allo-LCA, 3-oxo-DCA, 3-oxo-UDCA and t7k-LCA down-regulated the expression of both IL-17a and IL17f (Fig. 1l upper panel), with allo-LCA and 3-oxo-DCA being the two most active compounds. In addition, exposure of Treg cells specifically to CA, CDCA, 3-oxo- DCA, allo-LCA, and iso-allo-LCA increased the expression of FOXP3 and IL-10 (Fig. 1i, lower panel).

### Modulation of the fecal bile acid pool in murine models of IBD

To establish whether changes of the fecal bile acid pool observed in IBD patients were mechanistically linked to colitis development, a strategy for bile acid manipulation in TNBS or DSS colitic mice has been developed. Similarly to IBD patients, the two-mouse model of colitis exhibited an increase in the fecal excretion of primary bile acids and an increase of the primary/secondary bile acids ratio (Fig. 2a-e). The fecal excretion of five oxo- and allo- bile acids, 3-oxo-UDCA, 3-oxo-DCA, 3-oxo-LCA, allo-LCA, and iso-allo-LCA, was dramatically reduced in both models (Fig. 2b). Our attention was then focused on these compounds, whose concentration was also found to be modulated in patients with IBD (Fig. 1a-f). Correlation studies between fecal bile acids content and disease severity, expressed as colitis disease activity index (CDAI), demonstrated that the fecal levels of secondary bile acids correlated positively with the body weight and inversely with CDAI in both models (Fig. 2f). On the other hands, the primary/secondary ratio inversely correlated with the body weight in TNBS colitis model (Fig. 2f).

**Fig. 2.**
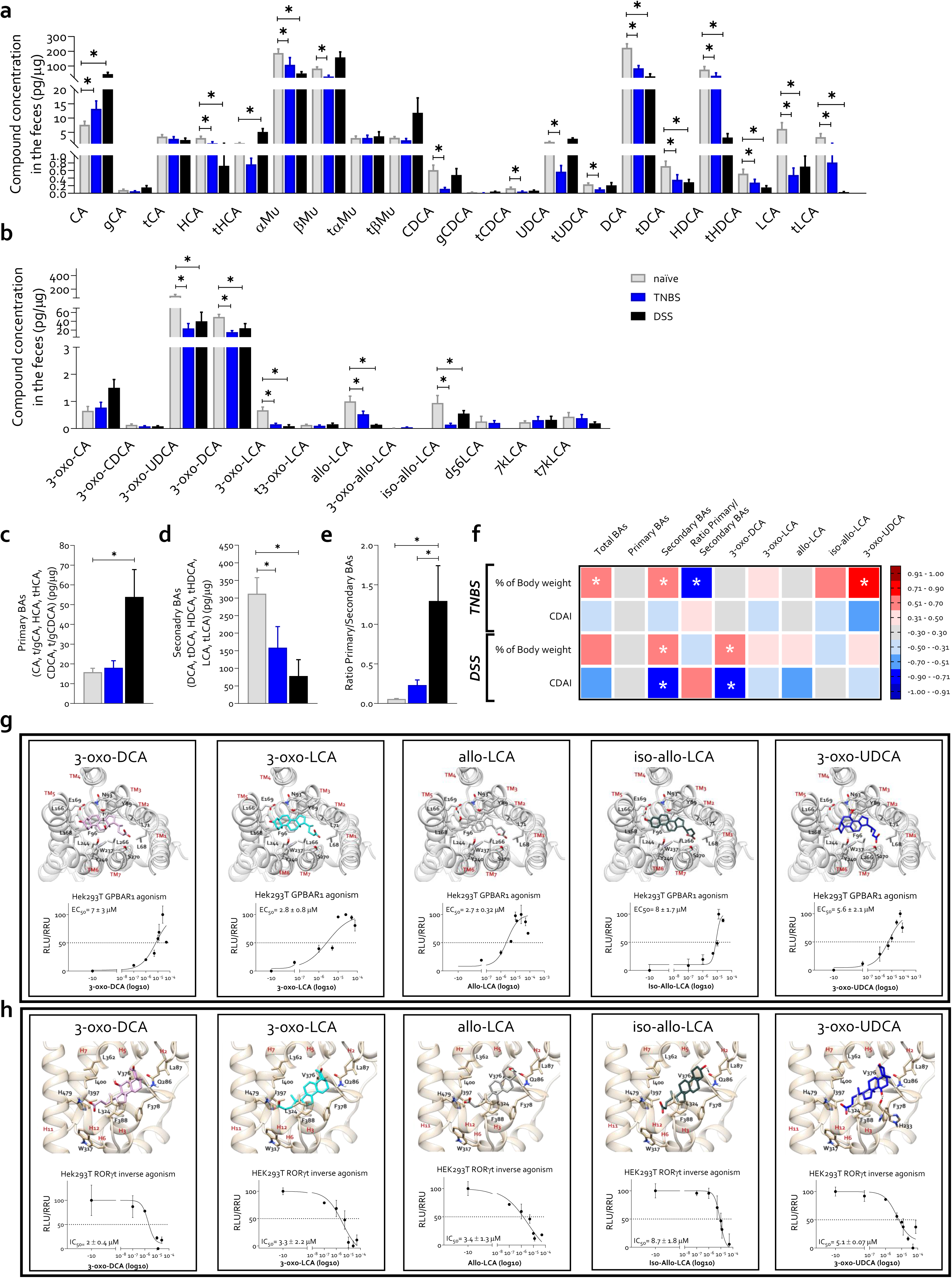
Fecal bile acid changes in mouse models of colitis. C57BL/6 male mice were treated with 2% of DSS in drinking water for 9 days or with 1 mg of TNBS via catheter into the intestinal lumen as described in Material and Method section. Fecal bile acid concentrations at the end of the experiments: (**a**) primary, secondary and conjugated bile acids, (**b**) oxo- and allo-derivatives of bile acids, (**c**) sum of all primary bile acids, (**d**) sum of all secondary bile acids and (**e**) ratio of primary to secondary bile acids. Data shown are the mean ± SEM of 5-12 mice per group. Statistical significance was assessed by 1-way ANOVA *p ≤ 0.05. (**f**) Correlation matrix between fecal bile acid concentrations and the clinical disease index measured using percentage of body weight and CDAI at the end of the experiments. Oxo- and allo- bile acid derivatives binding mode and potency evaluated respectively through docking calculations and transactivation assay on (**g**) GPBAR1 and (**h**) RORIZt receptors. The ligands are represented as sticks, while the receptors are represented as gray (GPBAR1) or tan (RORγt) ribbons with the interacting residues and helices displayed and labeled. Oxygen atoms are depicted in red, and nitrogen atoms are depicted in blue. H-bonds are displayed as dashed lines. Abbreviations: BAs, bile acids; TNBS, 2,4,6- trinitrobenzenesulfonic acid; DSS, Dextran sulfate sodium; T, tauro; LCA, lithocholic acid; DCA, deoxycholic acid; CDCA, chenodeoxycholic acid; HDCA, hyodeoxycholic acid; UDCA, ursodeoxycholic acid; HCA, hyocholic acid; αMu, alpha-muricholic acid; βMu, beta-muricholic acid; 7k, 7keto; CA, cholic acid; CDAI, colitis disease activity index; GPBAR1, G protein-coupled bile acid receptor; RORγt, RAR-related orphan receptor gamma t.

Particularly, fecal content of 3-oxo-UDCA showed a robust positive correlation with body weight changes and an inverse correlation with the CDAI in TNBS-colitis. Moreover, 3-oxo-DCA showed a direct correlation with the body weight and an inverse correlation with CDAI in the DSS-colitis model suggesting their potential involvement in disease development (Fig. 2f).

Since bile acids function as endogenous ligands for various membrane and nuclear receptors^15^, we have then investigated whether the oxo- and allo-derivatives exerted similar effects on these receptors (Fig. 2g, h and Table 1). The results demonstrated that none of the oxo- and allo- derivatives acted as an FXR or Vitamin D or Aryl Hydrocarbon Receptor (VDR and AHR) agonists (Table 1). However, all five bile acids acted as GPBAR1 agonists, with an EC_50_ of approx. 10 µM (Fig. 2g) and RORγt inverse agonists with an IC_50_ of 1-8 µM (Fig. 2h), thus defining the first dual GPBAR1 agonists and RORγt inverse agonists ever identified in mammals (Fig. 2g, h and Table 1). Docking calculation revealed that the five bile acids derivatives bind to both GPBAR1 and RORγt, adopting poses resembling those reported for GPBAR1 agonists and RORγt inverse agonist^28,29^. In GPBAR1, they interact with the key residues Y89^3.29^, Asn93^3.33^, Phe96^3.36^ and Trp237^6.48^ ^30^ critical for receptor activation. In RORγt, the derivatives carboxylic groups face His479 on helix H11, a residue crucial for stabilizing the close and active conformation of the receptor^31^.

### 3-oxo-DCA and allo-LCA counteract colitis development in a mouse model of TNBS-induced colitis

Since the intestinal content of oxo- and allo- bile acids derivatives were reduced in both IBD patients and models of colitis, we have then investigated whether supplementation of these bile acids species effectively modulates colitis development in murine colitis (Fig. 3-5 and fig. S2-S4). Administering C57BL/6J mice with TNBS promoted the development of severe disease with a mean weight loss of 20%, a CDAI of 9 and resulted in nearly 50% mortality (Fig. 3a-c). Treating these mice with the 3-oxo-DCA and allo-LCA, both at 10 mg/kg/day, attenuated disease severity and reduced colonic damage, as measured by assessing the colonic length and the number of leukocytes in the colon *lamina propria* (Fig. 3a-f). In contrast, 3-oxo-LCA failed to show any beneficial effect, while the 3-oxo-UDCA worsened disease severity and increased mortality, and the iso-allo-LCA exhibited intermediate levels of activity. The pathological findings were confirmed by the analysis of the intestinal expression of e-cadherin, a marker of mucosal integrity (fig. S3)^32^. The assessment of e-cadherin expression by immunohistochemistry demonstrated that while development of TNBS colitis was associated with a reduction of e-cadherin protein expression, this negative regulation was reversed by treating the mice with 3-oxo-DCA and allo- LCA but not by the other bile acids (fig. S2b).

**Fig. 3.**
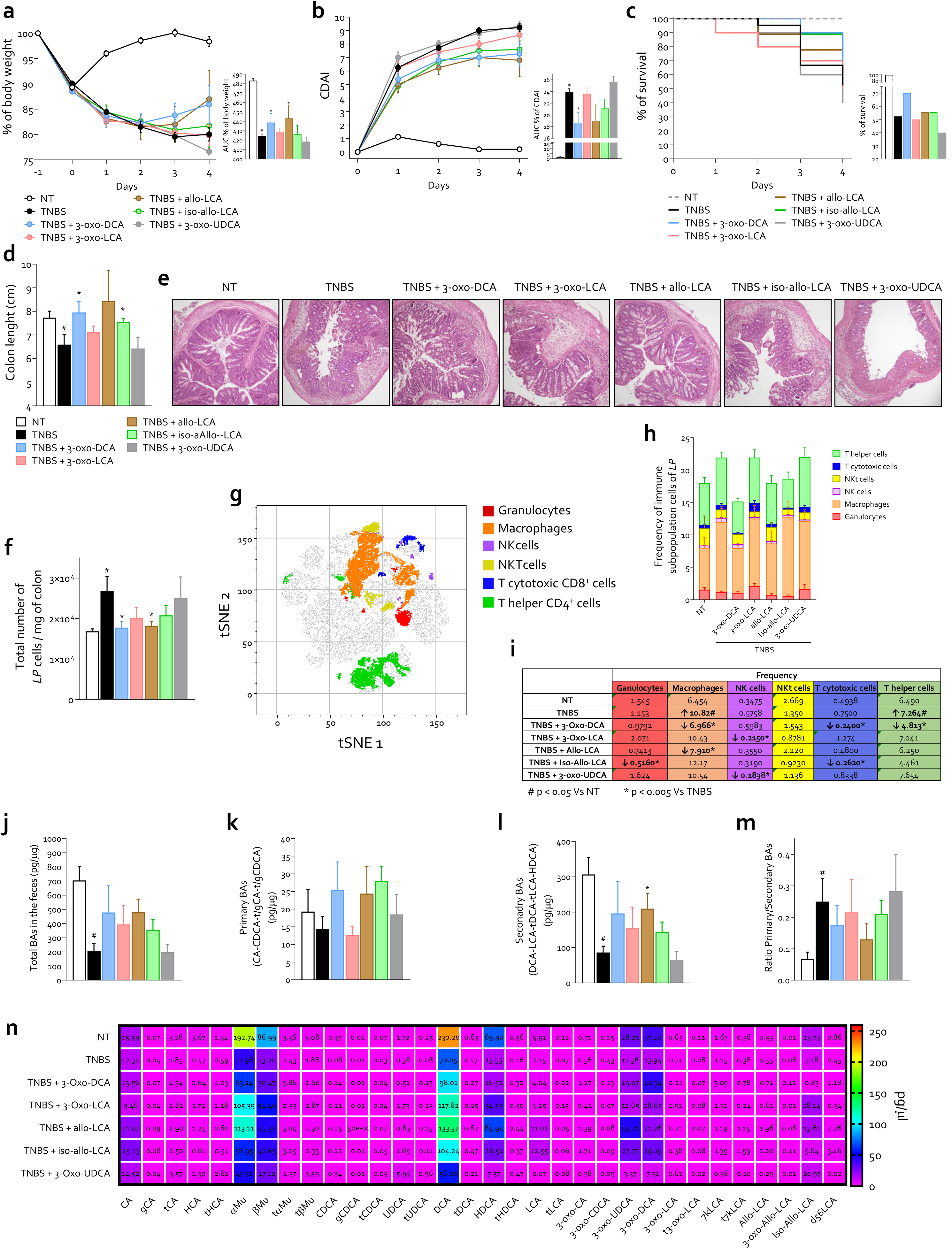
Modulation of colitis severity and leukocyte infiltration by oxo- and allo- derivatives of bile acids supplementation. C57BL/6 male mice were treated with TNBS and oxo- or allo- bile acids derivates: 3-oxo-DCA, 3-oxo-LCA, allo -LCA, iso-allo-LCA, 3-oxo-UDCA (10 mg/Kg/daily). Data shown are: (**a**) changes in body weight with area under the curve (AUC), (**b**) Colitis Disease Activity Index (CDAI) with AUC, (**c**) percent of survival of mice during the course of TNBS-induced colitis, (**d**) colon length and (**e**) hematoxylin and eosin (H&E) staining of colon (magnification 10×). (**f**) In order to establish the degree of inflammation the number of total leukocytes in the lamina propria was measured and then characterized with (**g-i** ) t-Distributed stochastic neighbor embedding (tSNE) analysis (**g-i** ). (**g**) Analysis in a t-SNE 2-dimensional (2D) scatterplot for leukocytes identified 6 subpopulations. (**h-i**) Frequency of leukocytes subpopulations in various experimental groups. (**j-n**) Analysis of bile acids in feces: (**j**) total bile acids pool in the feces (pg/μg), (**k**) primary bile acids (pg/μg), (**l**) secondary bile acids (pg/μg), (**m**) ratio between primary and secondary bile acids and (**n**) individual concentrations of bile acids and their derivates (pg/μg). Data shown are the mean ± SEM of 5-8 mice per group. Statistical significance was assessed by 1- way ANOVA *p ≤ 0.05. Abbreviations: NT, not treated; AUC, area under the curve; CDAI, colitis disease activity index; TNBS, 2,4,6-trinitrobenzenesulfonic acid; LP, lamina propria; tSNE, t-distributed stochastic neighbor embedding; NK, natural killer; NKT, natural killer T; BAs, bile acids; T, tauro; LCA, lithocholic acid; DCA, deoxycholic acid; CDCA, chenodeoxycholic acid; HDCA, hyodeoxycholic acid; UDCA, ursodeoxycholic acid; HCA, hyocholic acid; αMu, alpha-muricholic acid; βMu, beta-muricholic acid; 7k, 7keto; CA, cholic acid.

Furthermore, tSNE analysis allowed a characterization of leukocytes derived from colonic *lamina propria* (Fig. 3f-i). The results demonstrated that the development of TNBS colitis was associate with an increased inflow of macrophages and T helper cells (Fig. 3h, i). In contrast, the administration of 3-oxo-DCA effectively reduced the percentage of macrophages, helper T cells and cytotoxic T cells. Similarly, allo-LCA reduced only the percentage of macrophages and iso-allo-LCA reduced both the cytotoxic T cells and granulocytes. On the other hand, both 3-oxo-UDCA and 3-oxo-LCA did not exerted any effect on the total number of immune cells and reduced only the percentage of NK cells.

As shown in Fig. 3j-n and fig. S3, the TNBS administration in mice led to a decrease in the fecal content of total and secondary bile acids, including DCA, HDCA, 3-oxo-UDCA, 3-oxo-DCA, iso-allo-LCA and α/βMu (Fig. 3n). These changes were associated to the development of severe dysbiosis (Fig.4, r and fig. S4), as demonstrated by a strict reduction in the relative abundance of *Firmicutes* and an increase of *Bacteroidetes* and *Proteobacteria* (Fig. 4a-c)^33,34^, and consequent decrease in the Firmicutes/Bacteroidetes ratio^35^. Moreover, the correlation analysis demonstrated that while relative abundance of *Firmicutes* negatively correlated with disease severity, an increase of *Proteobacteria* worsened the disease outcome and positively correlated with the CDAI, confirming an opposite role of these two *phyla* in the development of intestinal inflammation (Fig. 4b, c).Furthermore, *Tenericutes* , whose relative abundance did not statistically change with disease induction, exhibited a strict direct correlation with body weight while *Bacteroidetes* showed an inverse correlation with CDAI (Fig. 4c). The administration of 3-oxo-DCA and allo-LCA in TNBS mice counteracted the severity of dysbiosis and restored the microbiota composition (Fig. 4a and c). At the genus level, colitis development was associated with a striking decrease in the abundance of *Clostridium, Blautia, Eubacterium, Helicobacter and Candidatus Arthromitus* , while relative abundance of *Erysipelatoclostridium, Enterococcus, Lactobacillus, Escherichia* and *Bacteroides* was increased (Fig. 4d, e). These data were consistent with human data^36–38^ ^39^. The correlation analysis in TNBS samples of genera with the severity of the disease revealed an inverse correlation between the relative abundance of *Clostridium, Marvinbryantia, Eubacterium, Blautia and Mycoplasma* and disease severity, along with a direct correlation of the relative abundance of *Enterococcus, Escherichia and Bacteroides* and degree of illness (Fig. 4f). Among the genus correlated with disease severity, TNBS statistically modulated *Clostridium, Enterococcus, Eubacterium, Blautia, and Escherichia*compared to naive mice (Fig. 4g). In this context, and only 3- oxo-DCA or allo-LCA reversed the effect exerted by TNBS, iso-allo-LCA showed a weaker beneficial effect and 3-oxo-LCA or 3-oxo-UDCA did not exert any effect. The analysis at the family level confirmed the results obtained at higher taxonomic levels, remarking the beneficial effect exerted by 3-oxo-DCA and allo-LCA on dysbiosis (fig. S4).

**Fig. 4.**
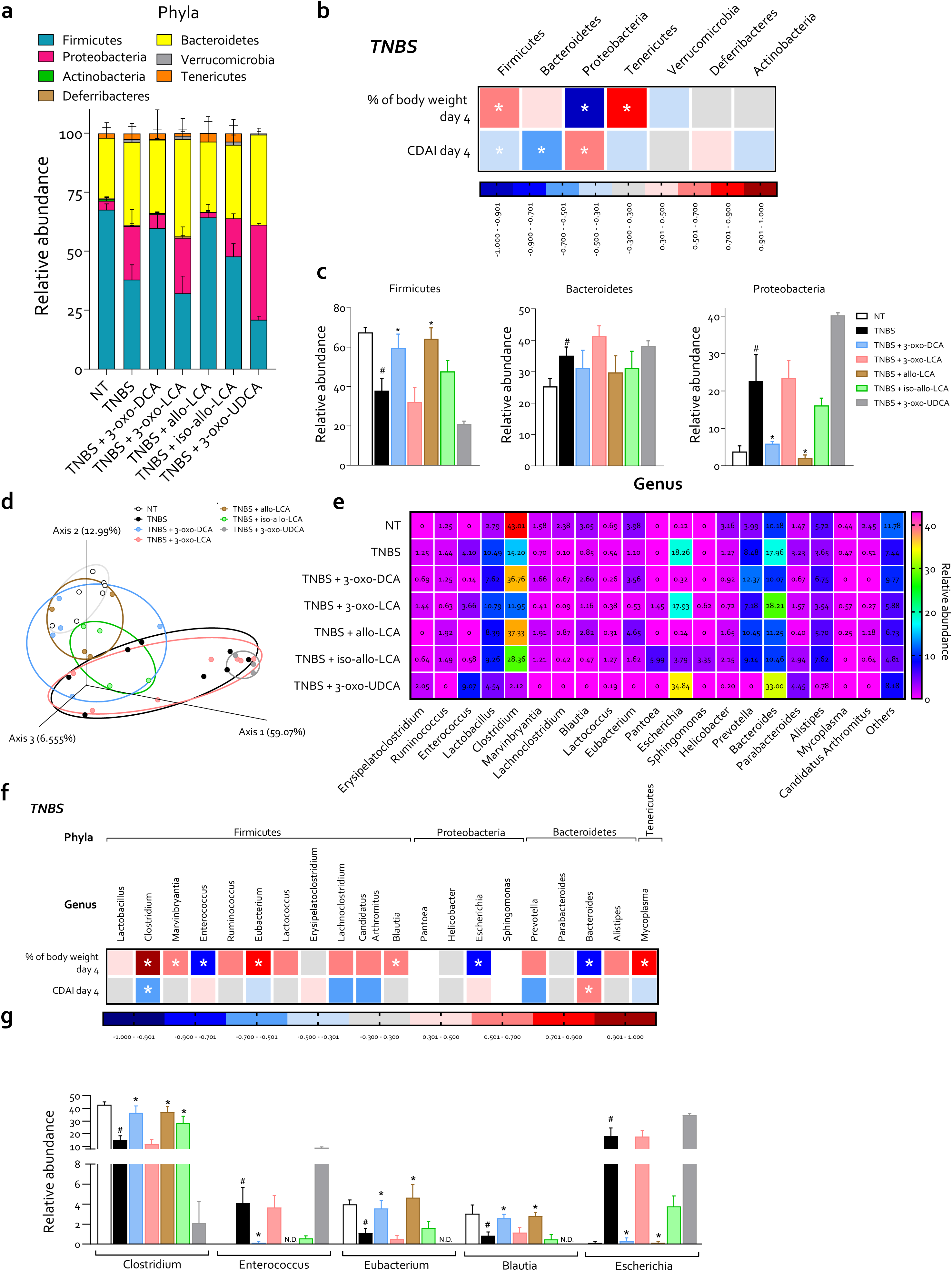
Reversal of dysbiosis by 3-oxo-DCA and allo-LCA in the mouse model of TNBS-induced colitis. C57BL/6 male mice were treated with TNBS and oxo- or allo- bile acids derivates: 3-oxo- DCA, 3-oxo-LCA, allo -LCA, iso-allo-LCA, 3-oxo-UDCA (10 mg/Kg/daily). (**a**) Relative abundance of phyla evaluated in fecal samples, (**b**) matrix correlation between relative abundance of phyla and percentage of weight loss and colitis disease activity index (CDAI) at the end of the treatment and in mice treated with TNBS alone and (**c**) representative histograms of phyla statistically modulated by colitis induction: Firmicutes, Bacteroidetes and Proteobacteria in each experimental group. (**d**) Analysis of the microbiota taxonomic profiles at the genus level using the principal component analysis (PCoA) plot of β diversity showing the distribution of different samples. (**e**) Relative abundance of genus identified in all experimental samples. (**f**) Correlation matrix between the relative abundance of genus and the severity of the disease, measured using percentage of body weight and CDAI. (**g**) Representative histograms of genus statistically modulated by colitis induction. Data shown are the mean ± SEM of 3-7 mice per group. Statistical significance was assessed by 1-way ANOVA *p ≤ 0.05. Abbreviations: NT, not treated; CDAI, colitis disease activity index; TNBS, 2,4,6-trinitrobenzenesulfonic acid; LCA, lithocholic acid; DCA, deoxycholic acid; UDCA, ursodeoxycholic acid.

The microbial taxonomic profiles at the species level are shown in Fig. 5 and table S1. The PCA diagram illustrate a robust separation of gut microbiota composition between the control and TNBS-treated groups and showed that only samples treated with 3-oxo-DCA and allo-LCA overlapped with control samples (Fig. 5a). By dissecting the statistically modulated species (Fig. 5b-g), we found that TNBS colitis associated with an expansion of *Bacteroides thataiotaomicron* ^40^*, Bacteroides acidifaciens, Enterococcus faecalis*^41–44^*, Enterococcus gallinarum*^45^*, Erysipellatoclostridium cocleatum*^46,47^*, Lactobacillus animalis, Lactobacillus murinus, Ruminococcus flavefaciens*^48^*, Escherichia coli*^45^*, Shigella flexneri*^49–51^ *and Akkermansia muciniphila*^52–54^ known to be related to pathogenesis of human IBD and experimental murine colitis. In addition, the development of TNBS colitis associated with a decreased abundance of *Clostidium sp* ^55^*, Clostridium fusiformis* , *Eubacterium plexicaudatum* ^56^ *and Marvinbryantia formatexigens*. The correlation analysis between the relative abundance of all species identified above 1% and the severity of the disease in TNBS samples revealed that 5 species positively correlated with disease severity, while 8 species exhibited an inverse correlation (Fig. 5h). Cross-referencing the data of species whose relative abundance was statistically altered by TNBS administration compared to naive mice (Fig. 5c-g) and species in TNBS-treated samples that correlated with disease severity, we were able to identify a community of bacterial species consisting of 8 species that correlate with the disease severity: *Bacteroides thetaiotaomicron, Bacteroides acidifaciens, Clostridium sp, Enterococcus faecalis, Enterococcus gallinarum, Marvinbryantia formatexigens, Escherichia coli and Shigella flexneri* . (Fig. 5h, species indicated in bold). Most bacteria that exhibited a positive correlation with disease severity showed a mild inverse correlation with fecal content of oxo- and allo- derivatives. Conversely, bacterial species showing negative correlation with disease severity, displaying a positive correlation with fecal content of 3-oxo-DCA (Fig. 5h). In support of this view, the fecal content of *Clostridium sp*, which harbors both the BSH and Bai genes, was decreased in diseased mice, dropping from 41% to 14%, while concomitant administration of 3-oxo-DCA reversed this change. In contrast, the relative abundance of *Bacteroides acidifaciens and Shigella flexneri*, both that lacking BSH, increased reaching a combined percentage of 12%, whereas they represented only 5% in naive mice ^57,58^. The *Bacteroides thetaiotaomicron, Enterococcus faecalis, and gallinarum* are known as BSH expressing species, but together they represented only 4%^57^. Thus, with the notable exception *Escherichia coli* , which is a BSH expressing species whose relative abundance rose to nearly 20%, these changes are consistent with view that the colitogenic metacommunity is characterized by a decreased expression of BSH^57,58^. These data highlight a mechanistic role of dysbiosis-induced deficiency of oxo- and allo- derivatives in promoting intestinal inflammation in rodent models of colitis.

**Fig. 5.**
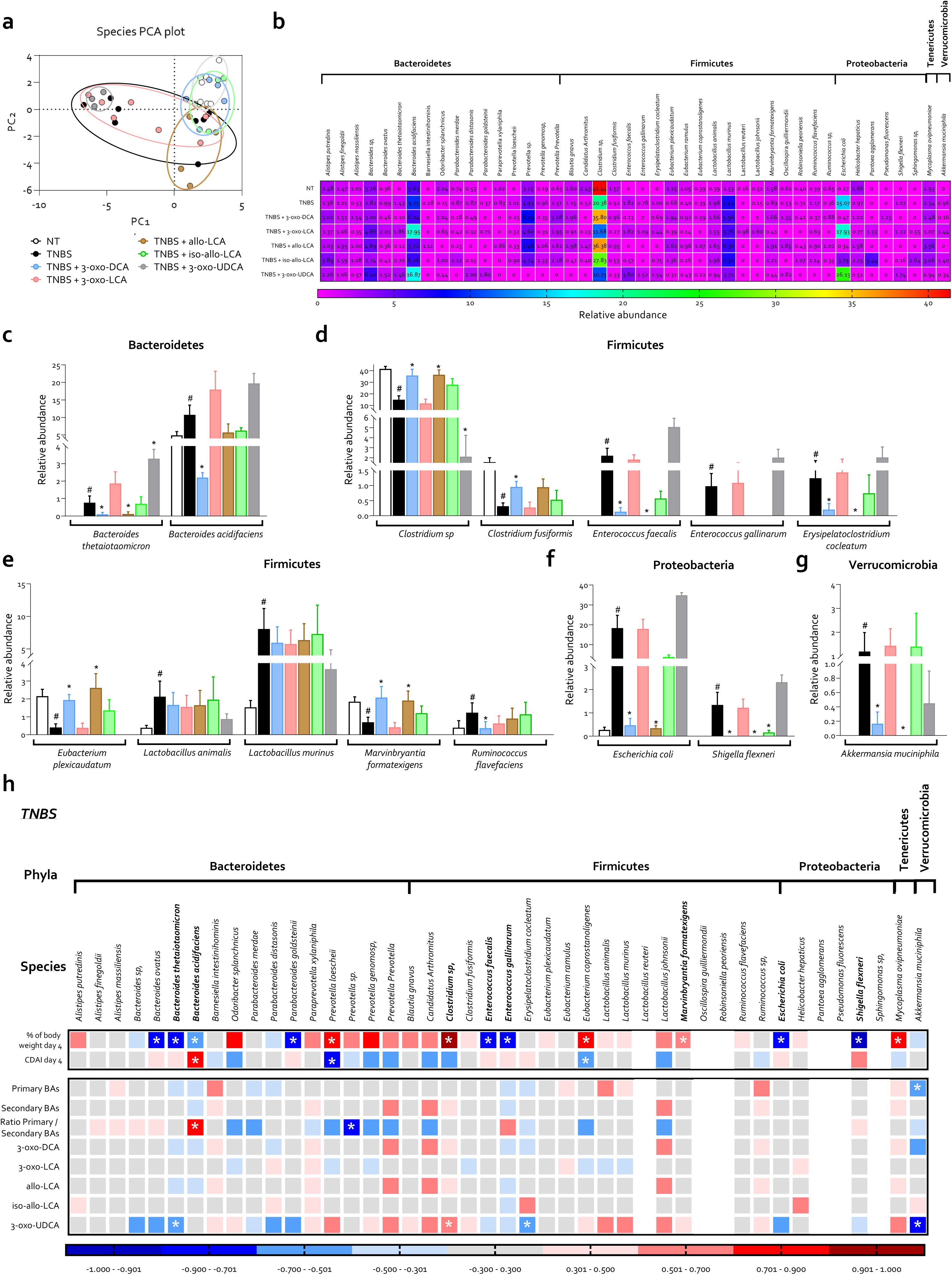
Expansion of pathogenic species in TNBS-treated mice and restoration by oxo- and allo- bile acid derivatives. C57BL/6 male mice were treated with TNBS and oxo- or allo- bile acids derivates: 3-oxo-DCA, 3-oxo-LCA, allo -LCA, iso-allo-LCA, 3-oxo-UDCA (10 mg/Kg/daily). (**a**) Analysis of the microbiota taxonomic profiles at the species level using the PCoA plot of β diversity showing the distribution of different samples. (**b**) Relative abundance of all species identified above 1% in experimental samples and (**c**-**g**) representative histograms of species statistically changed by colitis induction. (**h**) Correlation matrix between the relative abundance of all species identified above 1% and the severity of the disease, measured using percentage of body weight and CDAI and bile acid concentration in the feces, in TNBS samples. Species that showed significant correlations and were statistically modulated by colitis induction were indicated in bold. Data shown are the mean ± SEM of 3-7 mice per group. Statistical significance was assessed by 1-way ANOVA *p ≤ 0.05. Abbreviations: PCoA, principal component analysis; NT, not treated; TNBS, 2,4,6-trinitrobenzenesulfonic acid; BAs, bile acids; LCA, lithocholic acid; DCA, deoxycholic acid; UDCA, ursodeoxycholic acid.

In accordance with the data described above, 3-oxo-DCA and allo-LCA, which alleviated signs and symptoms of the disease (Fig. 3), modulate the relative abundance of identified species to restore eubiosis similar to that measured in naive mice (Fig. 5c-g).

### Anti-inflammatory effects of 3-oxo-DCA are partially GPBAR1-dependent

Since the data shown in Fig. 2g, h and Table 1 demonstrated that of 3-oxo-DCA is a dual GPBAR1 agonist and RORγt inverse agonist, we have dissected the role of these receptors by treating Gpbar1^+/+^ and Gpbar1^-/-^ mice rendered colitic by DSS and treated with 3-oxo-DCA (Fig. 6). The disease outcome, monitored by changes in body weight and CDAI, was worsened by Gpbar1 gene ablation, confirming the protective role of this receptor in the maintenance of intestinal homeostasis (Fig. 6a, b) ^18^. The administration of 3-oxo-DCA resulted in a very robust therapeutic effect in Gpbar1^+/+^ mice, and this effect was partially attenuated, although not completely reversed, in Gpbar1 ^-/-^ mice (Fig. 6a, b), as also confirmed by the pathology analysis (Fig. 6c, d).

**Fig. 6.**
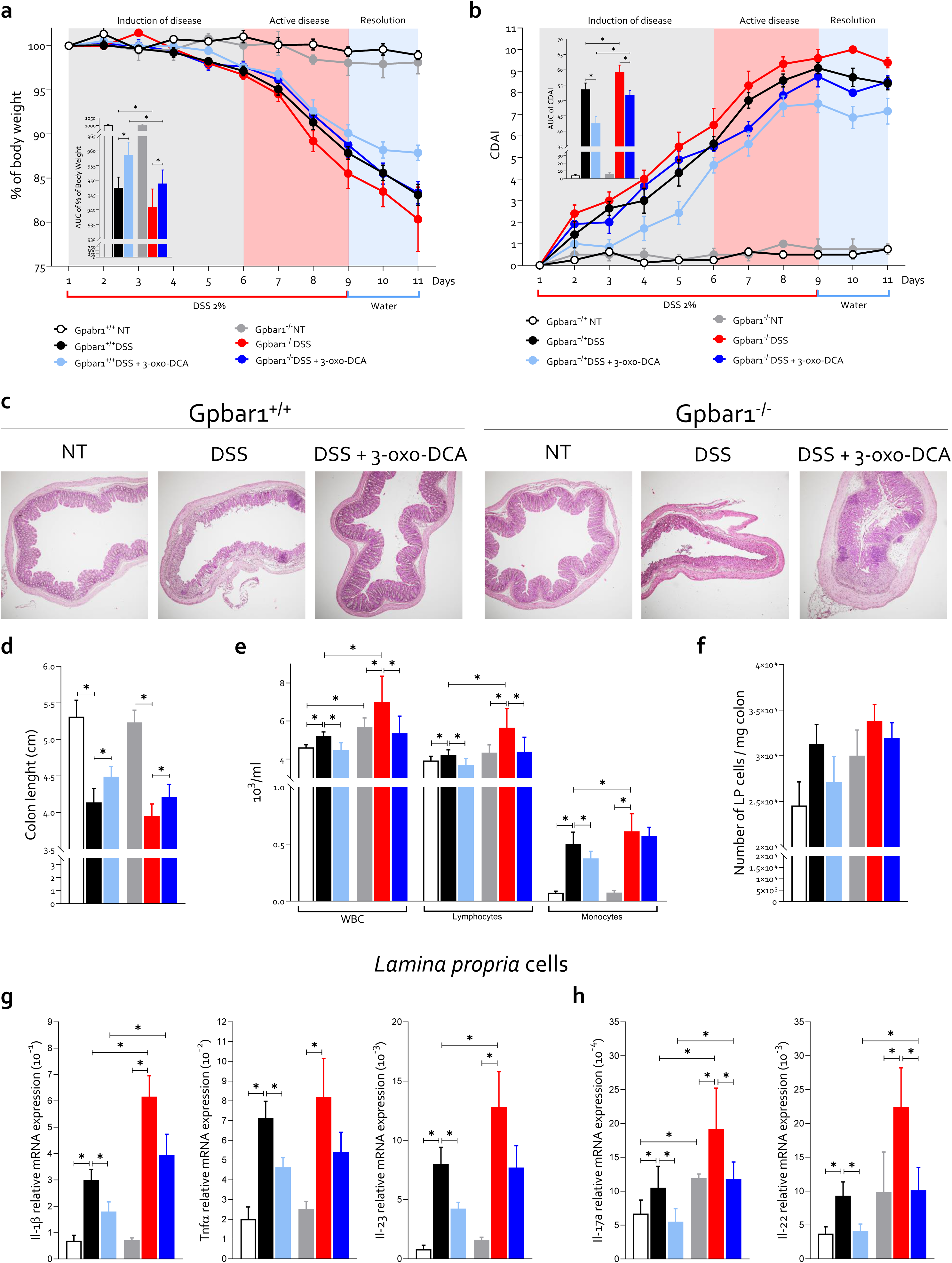
Effect of Gpbar1 ablation on 3-oxo-DCA treatment efficacy in the mouse model of DSS- induced colitis. Gpbar1^-/-^ C57BL/6 male mice and their wild types littermates were treated with DSS from day 1 to day 9 and from day 9 to day 11 with water. From day 1 to day 11, 3-oxo-DCA was administered via oral gavage at a dosage of 10 mg/kg/day. The severity of the disease was assessed by: (**a**) percentage of body weight change and area under the curve (AUC) of body weight trends, (**b**) Colitis Disease Activity Index (CDAI) and AUC of CDAI, (**c**) microscopic analysis of the colon using hematoxylin and eosin staining (magnification 10×), and (**d**) measurement of colon length. (**e**) Total number of white blood cells (WBC), lymphocytes and monocytes in the blood (10^3^ /ml) and (**h**) ratio between *lamina propria* cells and colon weight (mg). (**g-h** ) Relative mRNA expression of (**g**) Il-1β, Tnfα and Il-23 and (**h**) IL-17a and IL-22 measured in the *lamina propria* cells of the colon. Data are normalized to GAPDH mRNA. Data shown are the mean ± SEM of 4-12 mice per group. Statistical significance was assessed by 1-way ANOVA *p ≤ 0.05. Abbreviations: NT, not treated; DSS, Dextran sulfate sodium; DCA, deoxycholic acid; AUC, area under the curve; CDAI, colitis disease activity index; WBC, white blood cells; LP, lamina propria; Il- 1β, interleukin-1β; Tnfα, tumor necrosis factor-alpha; Il-23, interleukin-23; Il-17a, interleukin-17a; Il-22, interleukin-22.

Furthermore, administration of 3-oxo-DCA reduced the total number of WBC, as well as the number of lymphocytes and monocytes in the blood, and blunted the influx of immune cells in the colon (Fig. 6e, f). Naive Gpbar1^-/-^ mice exhibited a higher number of immune cells in the blood and in the *lamina propria* of the colon compared to wild-type mice, and treating these mice with DSS further increased these numbers (Fig. 6e, f). The protective effects exerted by 3-oxo-DCA on the number of circulating monocytes was largely abrogated by Gpbar1 gene ablation. In contrast, the therapeutic effects exerted by this agent on circulating lymphocytes were maintained demonstrating that 3-oxo-DCA exerts both GPBAR1-dependent and independent beneficial effects.

qRT-PCR analysis of signature cytokines produced by macrophages (Il-1β, TNFα, and IL- 23) and Th17 cells (Il-17a and Il-22) in the cells of *lamina propria* (Fig. 6g, h), revealed that disease induction led to increased production of pro-inflammatory cytokines in both murine strains. Administration of 3-oxo-DCA reversed intestinal inflammation in wild-type mice, while in Gpbar1^-/-^ mice, the beneficial effect of treatment was only partially maintained.

Since in contrast to GPBAR1, deletion of RORγt would result in colitis attenuation, we have then investigated whether pharmacological activation of RORγt with cintirorgon^59^ reversed/attenuated the therapeutic effects exerted by 3-oxo-DCA (fig. S5). As expected, treating DSS mice with cintirorgon worsened the disease severity, resulting in a severe weight loss and higher CDAI in comparison to mice treated with DSS alone (fig. S5a, b). In this context, administration of 3-oxo-DCA was still effective in reversing the severity of inflammation, although the efficacy was attenuated. To further understand the role of RORγt and 3-oxo-DCA in the various stages of the disease, mice were sacrificed on day 6, at the end of the induction phase, and on day 11, during the disease remission phase. Cintirorgon exacerbated the severity of colon inflammation by reducing colon length and increasing the number of infiltrating cells in the lamina propria at both time points (fig. S5c-e). The beneficial effect of 3-oxo-DCA was maintained even in mice treated with DSS + cintirorgon at both time points, as confirmed by macroscopic and microscopic analysis of the colon (fig. S5c-e) and also by measurement of colon mRNA expression of Tnfα and Il-23, (fig. S5f). Furthermore, treating mice with 3-oxo-DCA attenuated Th17 cell activation induced by DSS alone or in combination with cintirorgon, as assessed by measuring the mRNA expression of IL-17a and IL-22 at day 6 and 11 (fig. S5g).

### The administration of IL-23 exacerbates colitis in mice, but its effect is counteracted by the administration of 3-oxo-DCA

IL-23 is a pro-inflammatory cytokine that drives T cells differentiation toward the Th17 phenotype ^60–63^. The IL-23/IL-23R pathway plays a regulatory role in modulating the expression of RORγt in T cells and is crucially implicated in the progression of chronic inflammatory disorders, including IBD ^63–66^. This pathway serves as a validated target to assess the impact of RORγt in the protective effects exerted by 3-oxo-DCA. Consequently, our investigation delved into whether 3- oxo-DCA could rescue colitis induced by DSS and IL-23 (Fig. 7, 8 and fig. S7, 8).

**Fig. 7.**
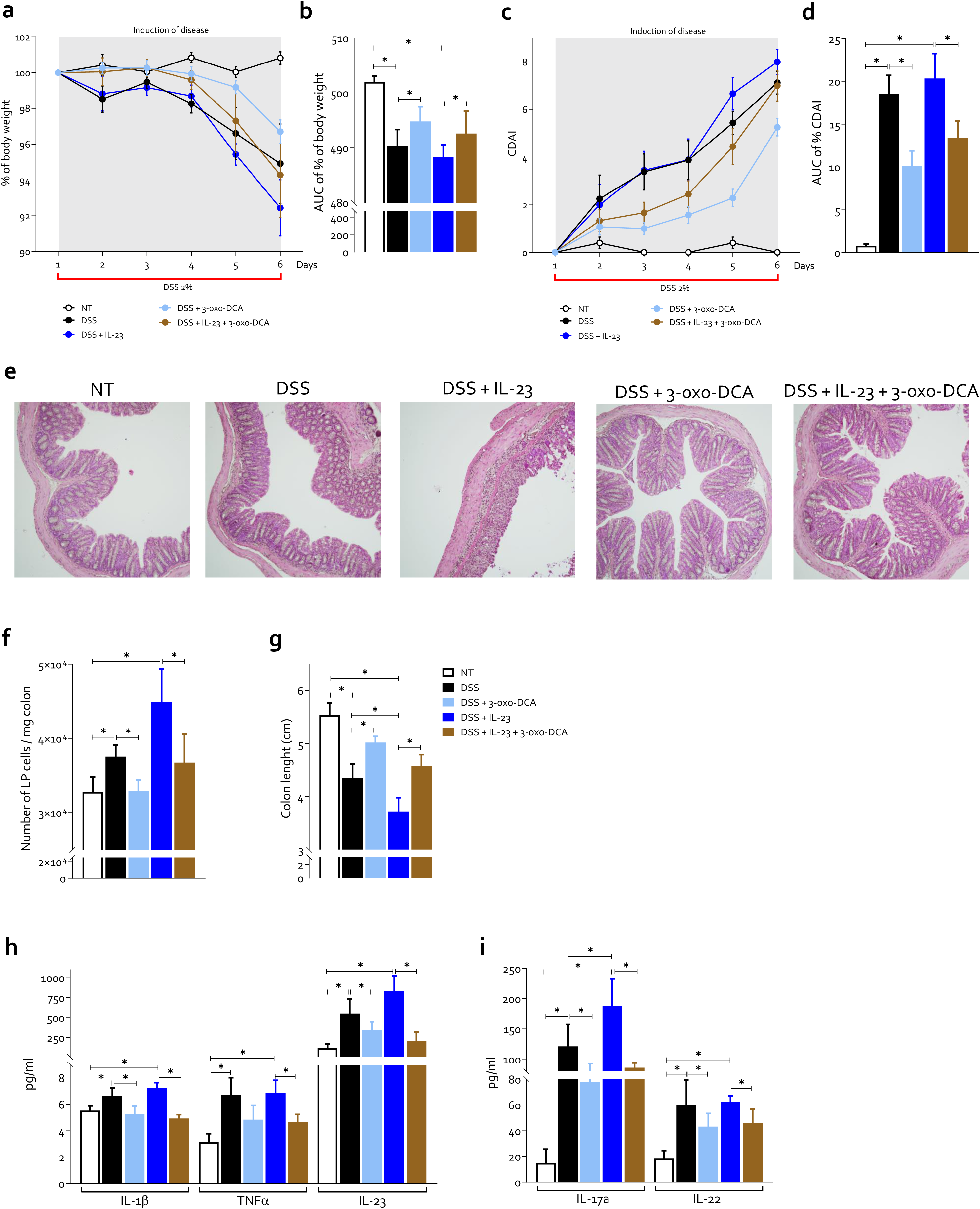
Impact of 3-oxo-DCA on IL-23-exacerbated DSS mouse model of colitis. Colitis was induced by simultaneous administration of DSS + IL-23 in C57BL/6 wild-type mice. 2% DSS was administered in drinking water for 6 consecutive days. IL-23 was administered at a dose of 500 ng/mouse via i.p. injection daily. 3-oxo-DCA (10 mg/Kg/daily) was administered by o.s. from day 1 to the end of the experiments. The mice were sacrificed at day 6. Disease severity was scored by the following evaluations: (**a**) changes in body weight (**b**) with area under the curve (AUC), (**c**) Colitis Disease Activity Index (CDAI), (**d**) with area under the curve (AUC), (**e**) hematoxylin and eosin staining of colon (magnification 10×), (**f**) ratio between *lamina propria* cells and colon weight (mg) and (**g**) colon length. (**h-i**) Concentration of (**h**) IL-1β, TNFα, IL-23 and (**i**) IL-17a and IL-22 in the serum of mice (pg/ml) evaluated by ELISA test at day 6. Data shown are the mean ± SEM of 4-12 mice per group. Statistical significance was assessed by 1-way ANOVA *p ≤ 0.05. Abbreviations: NT, not treated; DSS, Dextran sulfate sodium; DCA, deoxycholic acid; IL-23, interleukin-23; AUC, area under the curve; CDAI, colitis disease activity index; LP, lamina propria; Il-1β, interleukin-1β; Tnfα, tumor necrosis factor-alpha; Il-17a, interleukin-17a; Il-22, interleukin-22.

**Fig. 8.**
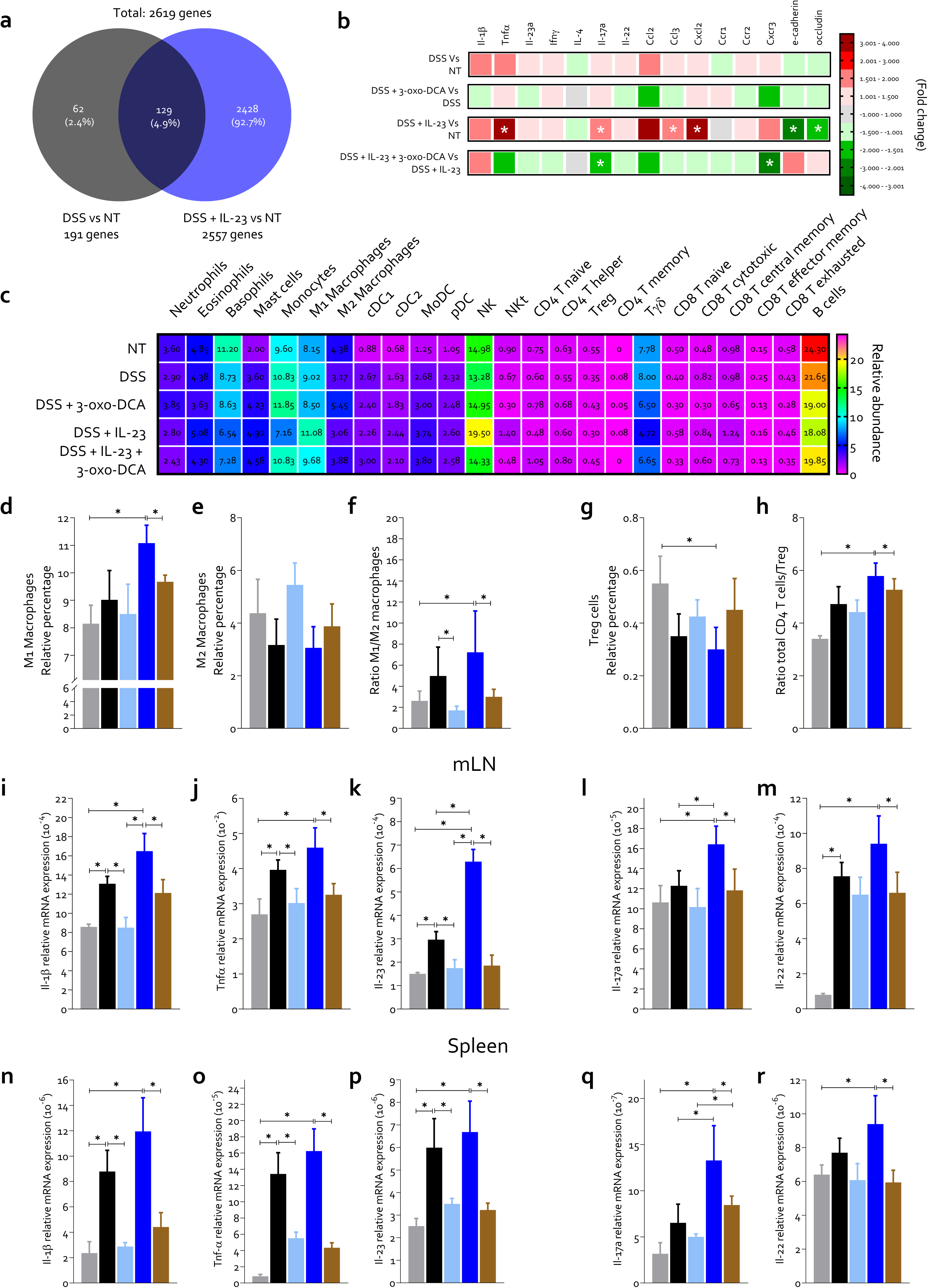
Gene expression changes and immune modulation in colonic tissue following disease induction and 3-oxo-DCA treatment. Colitis was induced by simultaneous administration of DSS + IL-23 in C57BL/6 wild-type mice. 2% DSS was administered in drinking water for 6 consecutive days. IL-23 was administered at a dose of 500 ng/mouse via i.p. injection daily. 3-oxo-DCA (10 mg/Kg/daily) was administered by o.s. from day 1 to the end of the experiments. The mice were sacrificed at day 6. (**a**) RNA-sequencing analysis was performed in colon samples and the Venn diagram of differentially expressed genes showing the overlapping regions between the mice experimental groups. (**b**) Some modulated genes belonging to inflammatory pathways. The up- regulated genes are indicated in green, while the down-regulated genes are identified in green. (**c-h**) Immune deconvolution of RNAseq data from colon of mice of different experimental group. Relative percentage of (**d**) macrophages M1, (**e**) M2 and (**f**) their ratio. Relative percentage of (**g**) Treg cells, (**h**) and ratio of total CD4 T cells and Treg. RNAseq data shown are the mean ± SEM of 4-6 mice per group. Statistical significance was assessed by 1-way ANOVA *p ≤ 0.05. (**i-m**) Relative mRNA expression of (**i**) Il-1β, (**j**) Tnfα, (**k**) Il-23, (**l**) Il-17a, and (**m**) Il-22 in mLN. (**n-r**) Relative mRNA expression of (**n**) Il-1β, (**o**) Tnfα, (**p**) Il-23, (**q**) Il-17a, and (**r**) Il-22 in spleen. qPCR data are normalized to GAPDH mRNA. Data shown are the mean ± SEM of 4-12 mice per group. Statistical significance was assessed by 1-way ANOVA *p ≤ 0.05. Abbreviations: NT, not treated; DSS, Dextran sulfate sodium; DCA, deoxycholic acid; IL-23, interleukin-23; Il-1β, interleukin-1β; Tnfα, tumor necrosis factor-alpha; Ifnγ, Interferon gamma; Il-4, interleukin-4; Il-17a, interleukin- 17a; Il-22, interleukin-22; Ccl2, CC Motif Chemokine Ligand 2; Ccl3, CC Motif Chemokine Ligand 3; Cxcl2, CXC Motif Chemokine Ligand 2; Ccr1, CC Motif Chemokine Receptor 1; Ccr2, CC Motif Chemokine Receptor 2; Cxcr3, CXC Motif Chemokine Receptor 3; cDC, conventional dendritic cells; MoDC, monocyte derived dendritic cells; pDC, plasmacytoid dendritic cells; NK, natural killer; NKt, natural killer t; Treg, regulatory T cells; mLN, mesenteric lymph nodes.

The clinical data, macroscopic and microscopic analysis of the colon revealed that the concurrent administration of DSS + IL-23 exacerbated the severity of colon disease (Fig. 7a-g). In addition, circulating levels of IL-1β, TNFα, IL-23, IL-17a, and IL-22 (Fig 7h, i) were robustly increased by treating mice with DSS, while the simultaneous administration of IL-23 increased only the production of IL-17a along with IL-22, confirming its promoting effect on Th17 cells. The 3-oxo-DCA alleviated the disease induced by DSS + IL-23, reducing all disease indices. 3-oxo-DCA reaffirmed its beneficial activity in colitis by reducing the concentration of all these cytokines, both in mice treated with DSS alone and in those treated with the combination of DSS/IL-23 (Fig. 7h, i).

To further investigate the mechanisms of the disease and the beneficial effect exerted by 3-oxo-DCA, we carried out RNA sequencing of the colon samples (Fig. 8a-h and fig. S6) and the analysis of the intestinal microbiota (fig. S7) of mice from each experimental group. The data demonstrated that at the day 6, the administration of DSS alone induced mild disease, modulating only 191 genes compared to untreated mice (Fig. 8a) resulting in only mild induction of expression of many cytokines, chemokines, and genes encoding tight junction proteins (Fig. 8b). In contrast, administration of DSS + IL-23 modulated 2557 genes compared to the untreated group, indicating a much more severe disease (Fig. 8a and fig. S6). Consistent with this view, Tnfα, Il-17a, Ccl3, and Cxcl2 were statistically up-regulated, while e-cadherin and occludin were down- regulated (Fig. 8b). The administration of 3-oxo-DCA modulated the expression of all these genes in the opposite manner compared to DSS and DSS + IL-23 (Fig. 8b). Immune deconvolution of RNAseq data revealed an increase in the percentage of pro-inflammatory macrophages with an elevated ratio of both M1/M2 macrophages and CD4 T cells/Treg (Fig. 8d-h). This inflammatory pattern was reversed by 3-oxo-DCA treatment in mice rendered colitic with DSS + IL-23, as well as in those administered with DSS alone. Intestinal microbiota analysis demonstrated that both DSS and DSS + IL-23 induced mild perturbations in the intestinal microbiota composition after 6 days of administration (fig. S7a). However, consistent with findings in the TNBS-induced colitis model (Fig. 4a-c), DSS and DSS + IL-23 administration decreased the abundance of *Firmicutes* while increasing *Bacteroidetes* and *Proteobacteria* inducing dysbiosis, also confirmed by the reduction in Shannon and Simpson index values (fig. S7b, c). Mild dysbiosis was associated with a modest alteration in the fecal bile acid pool with an increase in primary bile acids and a slight reduction in secondary bile acids (fig. S7d-f and fig. S8). 3-oxo-DCA exhibited a beneficial effect alleviating dysbiosis (fig. S8a-c) and changes in the fecal bile acid pool (fig. S7d-f and fig. S8).

To delve deeper into the modulation effect on the immune system exerted by disease induction and the administration of 3-oxo-DCA, we analyzed the expression of pro-inflammatory cytokines in the mesenteric lymph nodes (mLN) and spleen (Fig. 8i-r). The DSS administration up- regulated the expression of Il-1β, Tnfα, and Il-23 both in the mLN and spleen, confirming the involvement of macrophages in the immune response induced by colitis (Fig. 8i-k and n-p). Moreover, the up-regulation of the expression of the cytokines Il-17a and Il-22 produced confirmed the involvement of Th17 cells in the pathogenesis of the disease (Fig. 8l, m and q, r). This inflammatory pattern was significantly exacerbated by the simultaneous administration of DSS + IL-23. In contrast, the administration of 3-oxo-DCA down-regulated the expression of all these cytokines, alleviating the inflammation in both mLN and spleen (Fig. 8i-r).

### Microbiota-derived factors modulates immune polarization in colitis

The data presented so far demonstrated that colitis alters the intestinal microbiota in both IBD patients and murine models, leading to an alteration of the fecal content of bile acids and immune response (Fig. 1, 2). To investigate whether products of intestinal microbiota can alter immune cell polarization, we collected feces from mice treated for 9 days with DSS or DSS + 3- oxo-DCA, and homogenized and filtered them to remove all microorganisms. We then purified macrophages and T lymphocytes from the buffy coat of healthy donors and stimulated them with fecal homogenate (Fig. 9a). Analysis of M1 and M2 macrophages subpopulations through the expression of specific markers (CD38 and CD206, respectively), as well as analysis of TNFα and IL- 10 cytokine production by ELISA, demonstrated that fecal homogenate from DSS-treated mice exerted potent inflammatory activity, polarizing macrophages toward the M1 phenotype (Fig. 9b- e). Conversely, fecal homogenate from DSS + 3-oxo-DCA-treated mice significantly mitigated the pro-inflammatory effect, further upregulating CD206 and IL-10 expression compared (Fig. 9b-e). A similar effect was also observed in T lymphocytes (Fig. 9f-i). The fecal homogenate from DSS-treated mice induced an increase in RORC expression and IL-17 production, concurrently reducing FOXP3 expression and IL-10 production (Fig. 9f-i). Conversely, exposure of T lymphocytes to the fecal homogenate from DSS + 3-oxo-DCA-treated mice exerted an anti-inflammatory effect, attenuating the polarization towards the pro-inflammatory Th17 phenotype.

**Fig. 9.**
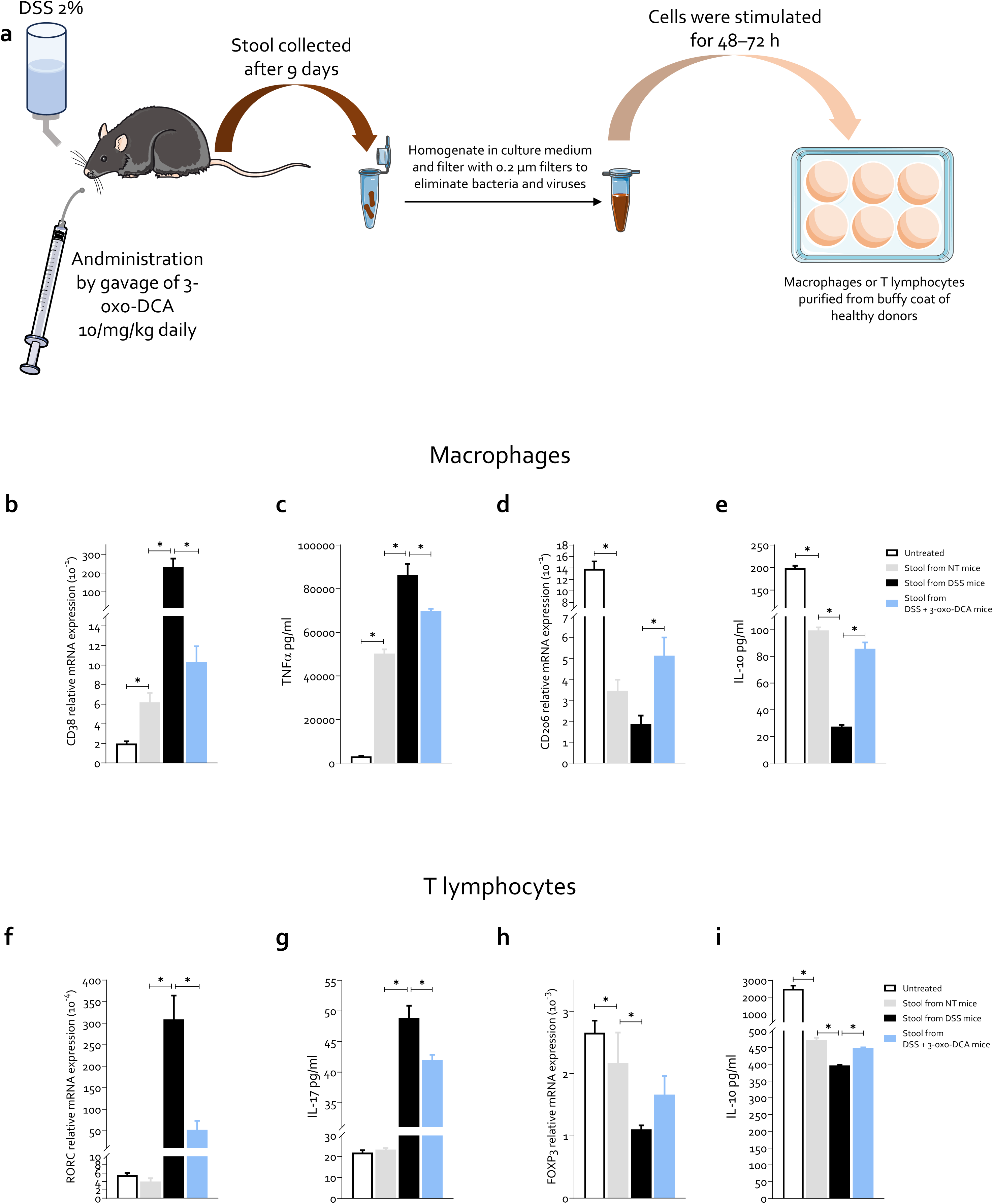
Influence of intestinal microbiota metabolites on immune cell phenotype. (**a**) Feces from mice treated with DSS or DSS + 3-oxo-DCA for 9 days were homogenized and filtered to remove all microorganisms. The obtained fecal homogenate was used to stimulate macrophages or T lymphocytes purified from the buffy coat of healthy donors. (**b-e**) The polarization of macrophages was measured by: (**b**) relative mRNA expression of the M1 marker CD38, (**c**) TNFα concentration in the cell culture medium, (**d**) relative mRNA expression of the M2 marker CD206 and (**e**) IL-10 concentration in the cell culture medium. (**f-i**) The polarization of T lymphocytes was measured by: (**f**) relative mRNA expression of the Th17 marker RORC, (**g**) IL-17 concentration in the cell culture medium, (**h**) relative mRNA expression of the Treg marker FOXP3 and (**e**) IL-10 concentration in the cell culture medium. qPCR data are normalized to GAPDH mRNA. Data shown are the mean ± SEM of 3-6 samples per group. Statistical significance was assessed by 1- way ANOVA *p ≤ 0.05. Abbreviations: NT, not treated; DSS, Dextran sulfate sodium; DCA, deoxycholic acid; CD38, cluster of differentiation 38; TNFα, tumor necrosis factor-alpha; CD206, cluster of differentiation 206; IL-10, interleukin-10; RORC, RAR Related Orphan Receptor C; IL-17, interleukin-17; FOXP3, Forkhead Box P3.

These data confirm that *in vivo* treatment with 3-oxo-DCA, likely by restoring eubiosis as shown in Fig. 4, 5, and fig. S4, 7, leads to the formation of a metabolome with anti-inflammatory characteristics that modulate the immune response toward a more tolerogenic phenotype.

## Discussion

IBD is a family of chronic disruptive disorder of the intestine that develops in genetically predisposed individual in response to a dysfunctional communication between a dysbiotic microbiota and the host immune system^2–5^. The intestinal microbiota generates a variety of chemical signals that are deemed essentially to maintain intestinal immune homeostasis and the perturbation of this network is increasingly recognized as the main mechanistic determinant in the development of IBD^67^. Bile acids are steroids generated at the interface of the host and bacterial metabolism^68^, in that, while primary bile acids, are synthesized in the liver from cholesterol, secondary bile acids represent the products of the intestinal microbiota. Despite various studies have revealed that these secondary bile acids exert a variety of regulatory effects on both the innate and adaptative immune system in the intestine^12^, the characterization of these bioactive steroids remains incomplete. Here, we report that 3-oxo-DCA, along with other oxo- and allo- derivatives of DCA, LCA and UDCA, define a novel class of dual GPBAR1 agonists/RORγt inverse agonists that promotes development of a tolerogenic phenotype by skewing macrophages and T helper cells polarization toward an M2 phenotype and Treg differentiation in IBD patients and mouse model of colitis. Supporting this view, we have found that fecal excretion of 3-oxo-DCA, along with other four oxo- and allo-DCA, LCA and UDCA derivatives, was selectively and significantly reduced in IBD patients and that this signature is reproduced in murine models of colitis. A detailed analysis of more than 30 bile acids in fecal samples demonstrated that the development of intestinal inflammation in IBD patients and model of colitis are associated with a decreased excretion of total bile acids which was almost completely sustained by a reduced excretion of secondary bile acids. Among the secondary bile acids, a statistically significant decrease occurred in five bile acids derivatives: 3-oxo-DCA, 3-oxo-LCA, allo-LCA, iso-allo-LCA, and 3-oxo-UDCA. This finding is consistent with previous published data that have shown that a reduction of secondary bile acids could be considered as a metabolomic signature for intestinal dysbiosis in IBD patients and animal models of colitis ^69–71^. Extending these observations, we now report that disease severity in CD patients inversely correlates with the relative abundance of 3-oxo-DCA in addition to DCA, while we have confirmed an inverse correlation between fecal levels of LCA and disease severity in UC patients (Fig. 1g). Consistent with this pattern, a reduction of secondary bile acids correlates with disease severity in both DSS and TNBS model of colitis. Notably, fecal levels of 3-oxo-DCA were robust predictor of intestinal inflammation in the DSS models, strengthening the mechanistic potential of this bile acid in the disease development (Fig. 2f). In addition, we have found that the disrupted pattern of bile acid excretion, namely a reduction of secondary bile acids, was associated with the development of an intestinal dysbiosis signed by a decrease, at the phylum level, of the *Firmicutes/Bacteroidetes* ratio^35^ and a significant increase of Proteobacteria in the mice models of colitis. Furthermore, examination at the species level, conducted in the mouse model allowed the identification of eight bacterial species that exhibited a direct correlation with disease severity and five species that showed an inverse correlation with the disease severity. By cross-referencing the data of species whose relative abundance underwent statistical alterations in the TNBS model, we have identified a microbial metacommunity formed by *Bacteroides thataiotaomicron, Bacteroides acidifaciens, Clostidium sp, Enterococcus faecalis, Enterococcus gallinarum, Marvinbryantia formatexigens, Escherichia coli and Shigella flexneri* whose relative abundance collectively predict likelihood to develop a severe disease in mice models of colitis. Moreover, the relative abundance of the members of this bacterial metacommunity was associated with the recruitment of inflammatory leukocytes, macrophages and T lymphocytes, in the colonic *lamina propria*, thereby indicating that the development of a colitogenic microbiota is the major determinant of leukocytes inflow in the inflamed intestine^72^. These findings are consistent with previous metagenomic studies showing that the relative abundance of microbial species capable of 7α/β- dehydroxylation of primary to secondary bile acids, effectively predicts clinical response to anti-cytokine or anti-integrin therapy in IBD patients^72^ . Together, these findings provide evidence that investigations of bile acids metabolome might represent a robust biomarker for disease severity and response to therapy in IBD.

To further elucidate the mechanistic potential of the above mentioned changes, we have investigated whether oxo- and allo- derivatives function as ligands for bile acid activated receptors^12^. By transactivation assay and docking studies, we were able to demonstrate that 3- oxo-DCA, 3-oxo-LCA, allo-LCA, iso-allo-LCA, and 3-oxo-UDCA effectively bind and transactivate GPBAR1 while function as inverse agonists of RORγt, with no activity on FXR, AHR, and VDR^73–75^ (Fig. 2g, h and Table 1). GPBAR1 is a GPCR for secondary bile acids^76^, highly expressed by myeloid cells^77,78^, and its activation suffices to rescue from inflammation in animal model of colitis^79^. Supporting the role for GPBAR1 in human IBD, the mutation analysis of the receptor has revealed a robust association of GPBAR1 single-nucleotide polymorphism rs11554825 with primary sclerosing cholangitis and UC^80^. In contrast to GPBAR1, whose expression is restricted to myeloid cells, RORγt is expressed by Th17 subsets and ILC3, and its activation prompts T cells differentiation toward Th17 proinflammatory phenotype, while reduces the number of intestinal Treg cells^75^.

Together these findings support the notion that 3-oxo-DCA, 3-oxo-LCA, allo-LCA, iso-allo- LCA, and 3-oxo-UDCA might represent a novel class of natural bile acids acting as dual GPBAR1 agonists/RORγt inverse agonists, with the potential to counter-regulate both the innate and adaptive immune response^17^. Confirming this concept, we have shown that challenging macrophages and T lymphocytes, purified from healthy donors, with 3-oxo-DCA, 3-oxo-LCA, allo-LCA, iso-allo-LCA and 3-oxo-UDCA skewed the macrophages polarization toward an M2, non-inflammatory phenotypes, and prevented Th17 polarization. Furthermore, treating TNBS colitis mice with 3-oxo-DCA protected against the development of clinical symptoms and histopathology features of colitis and reshaped the leukocytes trafficking, as shown by a dramatic reduction in the number of M1 macrophages and Th17 cells in the colonic lamina propria (Fig. 3). These beneficial effects extended to the composition of intestinal microbiota, since dosing colitic mice with 3-oxo-DCA protected against the development of the colitogenic bacterial metacommunity and reshaped bile acids metabolism^81^. Similar beneficial effects were obtained with allo-LCA, while the other bile acids exerted no effects or worsened, i.e. 3-oxo-UDCA, the disease severity.

Because the 3-oxo-DCA is a dual GPBAR1 and RORγt ligand, capable of modulating both the innate and adaptive immunity, we have then investigated how the two receptors contribute to the functional activity of this bile acid. For this purpose we induced colitis by administration of DSS to Gpbar1^-/-^ mice and wild type mice in which activation of RORγt was achieved by administering cintirorgon (a selective RORγT agonist^59^) (fig. S5) or IL-23 (a cytokine involved in the polarization of T helper cells towards the Th17 phenotype) (Fig. 7-8). The results of these studies demonstrated that genetic ablation of Gpbar1 or activation of the IL-23/IL-23R/RORγt/IL- 17 pathway reduced the efficacy of 3-oxo-DCA treatment, although it retained some of its beneficial effects. The partial loss of activity of 3-oxo-DCA in Gpbar1 ablated mice is consistent with the notion that GPBAR1 provides potent counter-regulatory signals to macrophages by inhibiting both NF-kB^82^ and NLPR3 inflammasome^83^ signaling and taking into consideration that Gpbar1 ablation reverses protection afforderd by LCA, a secondary bile acid, on both dysbiosis and immune dysfunction in the DSS model of colitis ^69^.

In addition to targeting the innate system, 3-oxo-DCA also reversed T cell activation as demonstrated by the detailed analysis of the immune response in the DSS/IL-23 model shown in Fig. 8. In this model, although treating DSS mice with IL-23 skewed the T cell polarization toward a Th17 phenotype, 3-oxo-DCA prevented these changes, as highlighted by histopathology analysis, immune deconvolution of RNAseq analysis and quantification of signature cytokines (Il- 1β, TNFα, Il-22 for monocytes/macrophages and Il-23 and Il-17a for Th17 cells), in the serum, in mLN and spleen (Fig. 8). Furthermore, evidence obtained *in vitro* on human macrophages and T lymphocytes challenged with bacterium-free homogenated of feces retrived from mice treated with DSS or with DSS + 3-oxo-DCA demonstrated that administration of 3-oxo-DCA *in vivo* promotes the development of tolerogenic signals that suffice to reverse M1 and Th17 differentiation in a bacterium free manner. (Fig. 9). In aggregate, these findings support the notion that colitis development associates with a reduction of 3-oxo-DCA while its restitution suffices to restore microbial/host immune homeostasis in models of colitis.

In conclusion, we have provided evidence that development of a dysbiotic microbiota in IBD and model of colitis impairs the production of secondary bile acids and selectively reduce the production of oxo- and allo-DCA, LCA and UDCA derivatives. The functional characterization of these bile acids, has lead to the discovey of 3-oxo-DCA as the first dual GPBAR1 agonist/RORγt inverse agonist that potently redirect the macrophages and Th17/Treg polarization. Recostitution of 3-oxo-DCA levels in mouse models of colitis reverses intestinal dysbiosis and inflammation in models of colitis highlighting a potential role for this agent in IBD management.

## Methods

### Human data

The metadata of patients, along with transcriptomic and metabolomic data, as well as microbiome profiling data derived from samples of healthy donors (HS, n = 26), UC patients (UC, n = 30), and Crohn’s disease patients (CD, n = 50), were retrieved from the Inflammatory Bowel Disease Multi’omics Database (IBDMDB)^71^.

CIBERSORTx^84^ was employed to infer the relative compositions of immune cells from the RNA-Seq data of the analyzed samples. The LM22 signature matrix, a validated leukocyte gene signature matrix consisting of 547 genes, was utilized to distinguish 22 human immune cell subsets. Following the estimation process (500 permutations), the inferred immune cell compositions were downloaded in .CSV format.

Correlation matrices were calculated using Spearman correlation to explore the relationships among bile acids, bacterial phyla, disease severity scores, gene expression, and immune cell populations.

### Purification of PBMCs from Buffy coat

Human peripheral blood mononuclear cells (PBMCs) were obtained from heparinized blood of voluntary healthy donors by density-gradient centrifugation using Ficoll-Hypaque and were cultured in RPMI-1640 medium containing 10% FBS for 6 hours.

### Human T lymphocytes and macrophages in vitro polarization

1 × 10^6^ of suspended lymphocytes were seeded into each well of 24-well treated by 200 μl PBS including 2 μg/ml anti-CD3 and 5 μg/ml anti-CD28 antibodies and were incubated with Th17-inducing cytokines (TGFβ, IL-6, IL1-β, IL-23, α-INFIT, α-IL4) or Treg-inducing cytokines (TGFβ, IL2), and bile acids (1 μM) for 72 h. Adherent monocytes were pre-differentiated into macrophages by culture for 6 days in RPMI/10% FCS supplemented with 50 ng/ml of either M-CSF or GM-CSF. Macrophages were polarized for 24 h to M1 macrophages using LPS (50 ng/ml) and IFNIT (20 ng/ml) and to M2 macrophages using IL-4, IL-10 and TGFβ (20 ng/ml) and treated with bile acids (1 μM).

In another experimental set, unpolarized macrophages and T lymphocytes were stimulated with a fecal homogenate for 24 and 72 hours, respectively. The feces were obtained from naive mice, treated with DSS alone or with DSS + 3-oxo-DCA for 9 days. Fecal samples were homogenized by adding 600 μl of culture medium per 100 mg of feces. To eliminate bacteria and viruses, the homogenates were filtered through 0.2 μm filters. For cell stimulation, 100 μl of homogenate per ml of medium were used.

### Luciferase reporter gene assay

Luciferase reporter gene assay protocol took place in 4 days. On day 1, HEPG2 or HEK293T were seeded in a 24-well plate at 7.5x104 cells/well or 2.5x104 cells/well respectively. Both lines were maintained at 37°C in humified atmosphere of 5% CO2. The next day cells were transiently transfected: separate mix for each receptor were prepared. FXR agonism: HepG2 cells were transfected with 200 ng of the reporter vector containing the response element p(hsp27) -TK-LUC, 100 ng of pSG5-FXR, 100 ng of pSG5-RXR, and 100 of pGL4.70 (Promega, Madison WI), a vector encoding the human Renilla gene. AhR agonism: HepG2 cells were transfected with 200 ng of the reporter vector pGL3-AhRE, 150 ng of pGL3-AhR and 150 ng of pGL4.70, the human Renilla gene. RORITt inverse agonism: HEK293T cells transfection mix was accomplished with 200 ng of the reporter vector pGL4.35, 150 ng of pFA-cmv-hRORC and 150 ng of pGL4.70, the human Renilla gene. GPBAR1 agonism: HEK293T cells were transfected with 200 ng of the reporter vector pG29- CRE-LUC, 150 ng of pcmvsport-hTGR5 and 150 ng of pGL4.70, the human Renilla gene. VDR agonism: HEK293T cells were transfected with 200 ng of the reporter vector pGL4.35, 150 ng of pFA-CMV-hVDR, and 150 of pGL4.70, the human Renilla gene. To promote transfection mixes were supplied by Fugene HD transfection reagent (Promega Madison WI). On the third day, cells were primed with agonist, CDCA 10 µM (FXR), Indirubine 1 µM (AhR), TLCA 10 µM (GPBAR1), LCA 10 µM (VDR) or bile acids 10 and 50 µM (for selected molecules further doses-response curve were set in a range of 0,1-75 µM). None agonist was necessary to promote RORITt activity, as it was considered to be constitutively expressed in naive cells. The last day, cellular lysate was assayed for luciferase and Renilla activities using the Dual-Luciferase Reporter assay system (Promega Madison WI). Luminescence was measured using Glomax 20/20 luminometer (Promega, Madison WI). Luciferase activities were normalized with Renilla activities.

### Docking

RORγt. The crystal structure of homo sapiens Retinoic acid receptor-related Orphan Receptor Gamma (RORγ) (PDB ID 3l0j) ^85^ was obtained from the Protein Data Bank website. The Nuclear receptor coactivator 2 (Src-2) in the RORγt active conformation, the co-crystallized ligands and water molecules were removed. Residues’ protonation states were assigned using the H^++^ webserver (http://newbiophysics.cs.vt.edu/H++/)^86–88^ at pH 7.4. The receptor was prepared using the Protein Preparation Wizard ^89^ tool implemented in Maestro ver. 11.8 using OPLS2005 forcefield ^90^ at pH 7.4.

GPBAR1. For GPBAR1, the homology model from D’Amore et al. ^91^ was used for docking calculations. The receptor underwent preparation steps as reported in Biagioli et al. ^92^

Ligands. The three-dimensional structures of allo-LCA, iso-allo-LCA, 2-oxo-DCA, 3-oxo- LCA and 3-oxo-UDCA were generated using the graphical user interface available in Maestro ver. 11.8.8 Protonation states at pH 7.4 in water was determined using the Epik module^93^ followed by a minimization process employing the OPLS 005 force field ^90^. This involved 2500 iterations utilizing the Polak-Ribiere Conjugate Gradient (PRCG) algorithm.

Docking Calculations. Glide software package^94^ was utilized for docking calculations, employing the Standard Precision (SP) algorithm of the GlideScore function^94,95^ and the OPLS2005 force field. Interaction grids, measuring 20 × 20 × 20 Å for RORγ and 25 × 16 × 17 Å for GPBAR1, centered on their respective ligand binding cavities, were generated. A total of 100 poses was generated, and ligand conformational sampling was increased two-fold compared to Glide’s default setting. The resulting binding poses were clustered based on their atomic RMSD, using a threshold of 2Å. Among five clusters, the conformation within the most populated cluster, with the lowest-energy values in both Glide Emodel and GlideScore, was considered

All figures were rendered by UCSF Chimera^96^.

### IBD mouse models

C57BL/6J wild-type mice were purchased from Envigo. GPBAR1 null mice (*Gpbar1*^-/-^) on C57BL/6 background were originally donated by Dr. Galya Vassileva (Schering-Plough Research Institute, Kenilworth). *Gpbar1*^-/-^ mice and their C57BL/6 congenic littermates were maintained in the animal facility of the University of Perugia under controlled temperature (22°C) and photoperiods (12:12-hour light/dark cycle), allowing unrestricted access to standard mouse chow and tap water.

We used four mouse models of chemically induced colitis to best simulate the broad spectrum of human IBDs and to better understand the mechanism of action of the tested compounds:

Colitis caused by trinitrobenzenesulfonic acid (TNBS) was induced in C57BL/6 wild-type mice. Briefly, mice were fasted for 1 day (day -1). On the following day (day 0), mice were anesthetized, and a 3.5 F catheter inserted into the colon such that the tip was 4 cm proximal to the anus. To induce colitis, 1 mg of TNBS (Sigma Chemical Co, St Louis, MO) in 50% ethanol was administered via catheter into the intestinal lumen using a 1 ml syringe (injection volume of 100 μl); control mice received 50% ethanol alone. When prompted by the experimental design, the 3- oxo-DCA, 3-oxo-LCA, Allo-LCA, Iso-allo-LCA or 3-oxo-UDCA (10 mg/Kg/daily) was administered by gavage (o.s.) from day 0 to the end of the experiments.

Colitis caused by DSS was induced in C57BL/6 wild-type mice as in Gpbar1^+/+^ and Gpbar1^-/-^ mice by administering 2% DSS (DSS: Dextran Sulfate, Sodium Salt of Affymetrix USA, molecular mass 40–50 kDa) in drinking water for 9 consecutive days. When prompted by the experimental design, the 3-oxo-DCA (10 mg/Kg/daily) was administered by gavage (o.s.) from day 1 to the end of the experiments.

To investigate the involvement of RORγt in the pathogenesis of IBD and in the mechanism of action of 3-oxo-DCA, we implemented a murine model of colitis induced by the administration of DSS + cintirorgon, a selective agonist of RORγt, in C57BL/6 wild-type mice. 2% DSS was administered in drinking water for 9 consecutive days. Cintirorgon was administered at a dose of 20 mg/kg via i.p. injection on day 1 and day 4. 3-oxo-DCA (10 mg/Kg/daily) was administered by gavage (o.s.) from day 1 to the end of the experiments. The mice were sacrificed at two time points: at the end of the disease induction phase (day 6) or during the resolution phase on day 11 (DSS administration was suspended on day 9).

In another murine experimental model, colitis was induced by simultaneous administration of DSS + IL-23 in C57BL/6 wild-type mice. 2% DSS was administered in drinking water for 6 consecutive days. IL-23 was administered at a dose of 500 ng/mouse via i.p. injection daily. 3-oxo-DCA (10 mg/Kg/daily) was administered by gavage (o.s.) from day 1 to the end of the experiments. The mice were sacrificed at the end of the disease induction phase (day 6).

The experimental protocols were approved by the Animal Care and Use Committee of the University of Perugia and by the Italian Minister of Health and Istituto Superiore di Sanità (Italy) and agreed with the European guidelines for use of experimental animals (permission n. 309- 2022-PR). The general health of the animals was monitored daily by the Veterinarian in the animal facility. On the day of sacrifice, mice were deeply anesthetized with sodium thiopental, 200 mg/kg b.w., and sacrificed before 12 P.M.

The severity of colitis was measured each day for each mouse by analyzing the body weight lost, the occult blood and stool consistency. Each parameter was scored from 0 to 4 and the sum represents the Colitis Disease Activity Index (CDAI). The scoring system had already been described in a previous work^97^.

### Histopathology

Colon sample (2-3 cm up anus) were first fixed in 10% Formalin, embedded in Paraffin, cut into 5-μm-thick sections and then stained with Hematoxylin/Eosin (H&E) for histopathological analysis.

Immunofluorescence was performed on paraffin embedded mouse colon. In brief, Ag retrieval was achieved on tissues sections for 90 min in the hot (95 °C) sodium citrate buffer (pH 6.0). Subsequently, slides were permeabilized with PBS and 0,1 % Triton and then incubated with Blocking buffer (PBS 1X with 10% horse serum and 1% BSA) for 1h at room temperature. Primary antibodies anti-E-CADHERIN (1:100) (GTX100443, Genetex 2456 Alton Pkwy Irvine, CA 92606 USA), were incubated overnight at 4 °C. The next day, after 3 washes with PBS 1X containing 0.1 % Tween 20 (PBST), sections were incubated with secondary antibody Alexa Fluor™ 568 A-11011 (1:1000) (Invitrogen, Thermofisher scientific Waltham, Massachusetts, USA), for 1h at room temperature in the dark. After 3 washes with PBST, nuclei were counterstained with DAPI 1X for 1 min in the dark and the reaction was stopped with a final wash in PBS 1X for 5 min. Then, slides were mounted with ProLong Glass Antifade Mountant (P36980) (Invitrogen, Carlsbad, CA), sealed with nail polish and observed at fluorescence microscope Olympus BX60. Immunofluorescence quantification was carried out as described in Shihan et al^98^

### Isolation of intestinal lamina propria cells

At the end of the experiments, the colons of mice were collected and cleaned of fecal contents. The cells were isolated from the colon *lamina propria* using the “Lamina Propria Dissociation Kit” from Miltenyi Biotec according to the manufacturer’s instructions.

### Flow-cytometry

Flow cytometry analyses on lamina propria cells were carried out using a three-laser configuration Thermo Fisher Scientific Attune Nxt flow cytometry system. Data were analyzed using FlowJo software (TreeStar). We analyzed the data with t-distributed Stochastic Neighbor Embedding (tSNE) using FlowJo software. The following monoclonal antibodies (mAbs) were used: CD45 APC (30-F11, Invitrogen); CD11b FITC (M1/70, BioLegend); Gr1 BV510 (RB6-8C5, BioLegend); CD3 PerCP-Cy5.5 (145-2C11, eBioscience); CD49b APC-eFluor780 (DX5, Invitrogen); CD4 SB436 (GK1.5 eBioscience); CD8 Super Bright 702 (53-6.7, Invitrogen).

### Bile acid determinations

Conjugated and un-conjugated bile acids concentrations of colon feces in mice experimental colitis models were measured by liquid chromatography-mass spectrometry (UHPLC-MRM-MS). Stock solutions and feces samples were prepared as reported in Biagioli et al.^99^.

UHPLC-MRM-MS analyses were performed both on a QTRAP 6500 (AB Sciex) equipped Shimadzu Nexera LC and Auto Sampler systems and on a Xevo TQS Micro (Waters) equipped with an Acquity UPLC H-class chromatographer. In both sets, BAs mixture was separated on a Luna Omega 1.6 μm Polar (C18, 100 Å, 50 × 2.1 mm; Phenomenex) at 40°C, and at a flow rate of 400 μl/min. For QTRAP 6500 set, the experimental conditions were reported in Biagioli et al.^99^. For Xevo TQS Micro set, the mobile phase A was H2O, 5 mM AmAc, 0.01% FA, and mobile phase B was MeOH/ACN (80:20), 5 mM AmAc, 0.01% FA. The gradient started at 50% B, increased to 55% B in 5 min and then to 70% B in 2 min and to 75% B in other 3 min, then was kept at 95% B for 5 min. The mass spectrometer was operated in negative MRM scanning mode using low collision energy to not fragment BAs, with the following parameters: cone potential at -50 V, collision energy at -5V and cell exit potential (CXP) at 20 V. Cone gas was set at 15 L/min, ion source gas at 300 L/min and ion spray voltage at -2500. Chromatograms were analyzed through the Quanlynx tool of the Masslynx software (Waters). 32 BAs have been identified and quantified thanks to the opportune standards curves as reported by Biagioli et al.^99^.

### RNA isolation and qRT-PCR

Total mRNA extraction from colon samples was performed using Tri-Reagent (Zymo Research) and Direct-zol™ RNA MiniPrep w/ Zymo-Spin™ IIC Columns (Zymo Research, Irvine, CA). After purification from genomic DNA using DNase I (Thermo Fisher Scientific, Waltham, MA), 1 μg of RNA from each sample was reverse transcribed using Kit FastGene Scriptase Basic (Nippon Genetics, Mariaweilerstraße, Düren, Germaniain) in 20-μl of reaction volume; 50 ng of cDNA was amplified in a 20-μl solution containing 200 nM each primer and 10 μl of PowerUp™ SYBR™ Green Master Mix (Thermo Fisher Scientific, Waltham, MA). All reactions were performed in triplicate using the following thermal cycling conditions: 2 min at 95°C, followed by 40 cycles of 95°C for 3 s, 60°C for 30 s, using a QuantStudio 3 system (Applied Biosystems, Foster City, CA). The relative mRNA expression was calculated accordingly to the ΔCt method. The primer used were as following (forward and reverse): mIl-1β: GCTGAAAGCTCTCCACCTCA and AGGCCACAGGTATTTTGTCG; mTnfα: GCCTCTTCTCATTCCTGCTT and GAGGCCATTTGGGAACTTCT; mIl-23: GACCCACAAGGACTCAAGGA and AGGCTCCCCTTTGAAGATGT; mIl-17a: TCCAGAAGGCCCTCAGACTA and TGAGCTTCCCAGATCACAGA; mIl-22: CAACTTCCAGCAGCCATACA and ATGAGCCGGACATCTGTGTT; hCD38: CCTGGCTGAAGTGACGTTATC and ACCTCCAGAGGTTGAGCAAA; hTNFα: AGCCCATGTTGTAGCAAACC and TGAGGTACAGGCCCTCTGAT; hCD206: GAGGAAAAGCTGCCAACAAC and CCAATCCAGAGTCCTGAGGT; hIL-17a: ACCAATCCCAAAAGGTCCTC and ACTTTGCCTCCCAGATCACA; hIL-17f: TGTCACGTAACATCGAGAGC and ACTGGGCCTGTACAACTTCC; hFOXP3: TCTTCCTTGAACCCCATGC and CCTGGAGGAGTGCCTGTAAG.

### Gut microbiota analysis

#### DNA extraction

The microbial DNA was purified from colon fecal samples of mice using the PureLink Microbiome DNA Purification Kit (Thermo Fisher Scientific, Waltham, MA), according the manufacturer’s instructions. The isolated DNA was quantified with a Qubit dsDNA HS Assay Kit on Qubit 3.0 Fluorometer (Thermo Fisher Scientific, Waltham, MA) according the manufacturer’s instructions and then stored at -20 °C.

#### 16S rDNA sequencing

Library and template preparation of the barcoded libraries were performed using Ion AmpliSeq™ Microbiome Health Research Kit (Thermo Fisher Scientific, Waltham, MA) and Ion 510 & Ion 520 & Ion 530 Kit - Chef on the Ion Chef platform (Thermo Fisher Scientific, Waltham, MA) according to the manufacturer’s instructions. Sequencing was performed on the Ion S5 platform and automated analysis, annotation, and taxonomical assignment were generated using Ion Reporter Software - AmpliSeq Microbiome Health Workflow (Ion Reporter 5.18.4.0). The Ion Reporter Software enables the rapid identification (at family, genus or species level) of microbes present in each sample, using Curated Greengenes v13.5 reference databases. The identification of families that produce 7αHSDH and BSH was made thanks to the literature data ^100–102^.

### AmpliSeq transcriptome

High-quality RNA was extracted from colon samples using Tri-Reagent (Zymo Research) and Direct-zol™ RNA MiniPrep w/ Zymo-Spin™ IIC Columns (Zymo Research, Irvine, CA) according to the manufacturer’s instructions. RNA quality and quantity were assessed with the Qubit® RNA HS Assay Kit and a Qubit 3.0 fluorometer followed by agarose gel electrophoresis. Libraries were generated using the Ion AmpliSeq™ Transcriptome Mouse Gene Expression Core Panel and Chef-Ready Kit (Thermo Fisher Scientific), according the manufacturer’s instructions. Barcoded libraries were combined to a final concentration of 100 pM, and used to prepare Template-Positive Ion Sphere™ (Thermo Fisher Scientific, Waltham, MA) particles to load on Ion 540™ Chips, using the Ion 540™ Kit-Chef (Thermo Fisher Scientific, Waltham, MA). Sequencing was performed on an Ion S5™ Sequencer with Torrent Suite™ Software v5.18 (Thermo Fisher Scientific). The analyses were performed with a range of fold <−2 and >+2 and a p value < 0.05, using Transcriptome Analysis Console Software (version 4.0.2), certified for AmpliSeq analysis (Thermo-Fisher).

### Multiplex Assay-Inflammatory Molecules

A multiplex biometric ELISA-based immunoassay was used according to the manufacturer’s instructions (MERCK cat. n. MTH17MAG-47K-05) using the Bio-Plex 200 instrument (Bio-Rad Laboratories). The following molecules were measured on mouse serum: IL- 1β, TNFα, IL-23, IL-17a and IL-22. Measurements were performed in duplicate. The analytes concentration was calculated using a standard curve, with software provided by the manufacturer (Bio-Plex Manager Software).

### Statistical analysis

We first performed the Kolmogorov-Smirnov test for normal distribution. The one-way ANOVA or unpaired Student t test were used for statistical comparisons (*p <0.05). The correlation analysis was performed using Pearson r in the case of a normal distribution of data, or Spearman r in the case of a non-normal distribution. All analyses were performed using the Prism 8.0 software (GraphPad).

### Data Availability Statement

The complete data of molecular docking calculations presented in this study are openly available in Mendeley data repository DOI: *10.17632/vdkn2x9g8n.1*. The complete UHPLC-MRM-MS data related to the fecal bile acids quantification that supporting the findings hereby presented are openly available in Mendeley data repository DOI: *10.17632/hmkvmsxrmb.1*. The complete RNA-seq data of mouse colon and the analysis of mouse intestinal microbiota presented this study are openly available in Mendeley data repository DOI: *10.17632/cdffdhvxnc.1*.

## Acknowledgements

This work was partially funded by PRIN-2022 (N. 20223K7L88).

## Author contributions

Mi.B. contributed conceptualization, methodology, investigation, visualization, funding acquisition, supervision, writing-original draft; C.D.G. contributed methodology and investigation; C.M. contributed methodology and investigation; S.M. contributed conceptualization, methodology, investigation, visualization, supervision and writing-review & editing; R.B. contributed methodology and investigation; Ma.B. contributed methodology and investigation; G.U. contributed methodology and investigation; R.R. contributed methodology and investigation; G.L. contributed methodology and investigation; R.S.U.K. contributed methodology and investigation; M.C.M. contributed conceptualization, methodology, investigation, and writing-review & editing; E.M. contributed methodology and investigation, F.D.P. contributed methodology and investigation; B.Ch. contributed methodology and investigation; B.F. contributed methodology and investigation; B.Ca. contributed conceptualization, supervision, and writing-review & editing; L.C. contributed methodology and investigation; G.N. contributed methodology and investigation; P.R. contributed conceptualization and writing-review & editing; E.D. contributed conceptualization and writing-review & editing; A.Z. contributed conceptualization, funding acquisition, project administration, supervision, writing-original draft and writing-review & editing; S.F. contributed conceptualization, funding acquisition, project administration, supervision, writing-original draft and writing-review & editing.

## Data and materials availability

All data are available in the main text or in the supplementary materials.

