## Supplementary Data for "A microbial derived bile acid acts as GPBAR1 agonist and RORγt inverse agonist and reverses inflammation in inflammatory bowel disease"

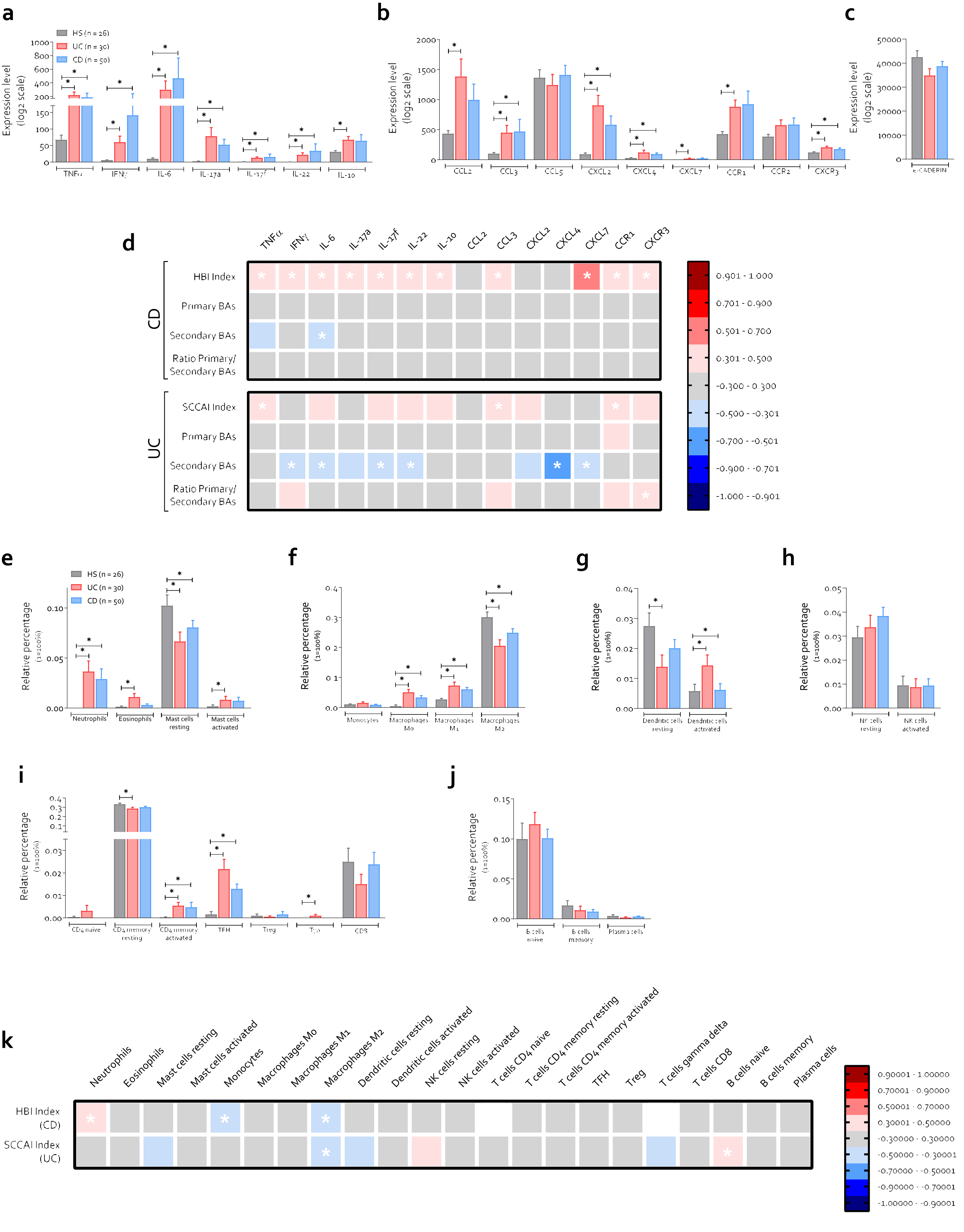
a
HS (n = 26)
UC (n = 30)
CD (n = 50)
Expression level (log2 scale)
TNFα
IFNγ
IL-6
IL-17a
IL-17f
IL-22
IL-10
b
Expression level (log2 scale)
CCL2
CCL3
CCL5
CXCL2
CXCL4
CXCL7
CCR1
CCR2
CXCR3
c
Expression level (log2 scale)
d
HBI Index
Primary BAs
Secondary BAs
Ratio Primary/ Secondary BAs
SCCAI Index
CD
UC
0.901 - 1.000
0.701 - 0.900
0.501 - 0.700
0.301 - 0.500
-0.300 - 0.300
-0.500 - -0.301
-0.700 - -0.501
-0.900 - -0.701
-1.000 - -0.901
e
Relative percentage (1=100%)
Neutrophils
Eosinophils
Mast cells resting
Mast cells activated
f
Monocytes
Macrophages M0
Macrophages M1
Macrophages M2
g
Dendritic cells resting
Dendritic cells activated
h
NK cells resting
NK cells activated
i
CD4 naive
CD4 memory resting
CD4 memory activated
TFH
Treg
CD8
j
B cells naive
B cells memory
Plasma cells
k
Neutrophils
Eosinophils
Mast cells resting
Mast cells activated
Monocytes
Macrophages M0
Macrophages M1
Macrophages M2
Dendritic cells resting
Dendritic cells activated
NK cells resting
NK cells activated
T cells CD4 naive
T cells CD4 memory resting
T cells CD4 memory activated
TFH
Treg
T cells gamma delta
T cells CD8
B cells naive
B cells memory
Plasma cells
HBI Index (CD)
SCCAI Index (UC)
0.90001 - 1.00000
0.70001 - 0.90000
0.50001 - 0.70000
0.30001 - 0.50000
-0.30000 - 0.30000
-0.50000 - -0.30001
-0.70000 - -0.50001
-0.90000 - -0.70001
-1.00000 - -0.90001

**Supplementary figure 1.** The analysis of an Inflammatory Bowel Disease (IBD) cohort from the Human Microbiome Project (HMP2). Data shown are: (**a**) cytokines, (**b**) chemokines and (**c**) e-caderin expression in the rectum of healthy individuals, patients with ulcerative colitis (UC), and patients with Crohn's disease (CD). (**d**) Correlation of disease severity indexes of UC and CD patients with cytokines and chemokines expression. Identification of (**e-j**) relative compositions of immune cells from the RNA-Seq data of the analyzed samples by a validated leukocyte gene signature matrix. (**k**) Correlation between disease severity indexes of UC and CD patients and immune cells populations. Abbreviations: HS, healthy samples; UC, ulcerative colitis; CD, Crohn's disease; TNFα, tumor necrosis factor-alpha; IFNγ, Interferon gamma; IL-6, interleukin-6; IL-17a, interleukin-17a; IL-17f, interleukin-17f; IL-22, interleukin-22; IL-10, interleukin-10; CCL2, CC Motif Chemokine Ligand 2; CCL3, CC Motif Chemokine Ligand 3; CCL5, CC Motif Chemokine Ligand 5;CXCL2, CXC Motif Chemokine Ligand 2; CXCL4, CXC Motif Chemokine Ligand 4; CXCL7, CXC Motif Chemokine Ligand 7; Ccr1, CC Motif Chemokine Receptor 1; Ccr2, CC Motif Chemokine Receptor 2; Cxcr3, CXC Motif Chemokine Receptor 3; HBI, Harvey-Bradshaw Index; SCCAI, Simple Clinical Colitis Activity Index.

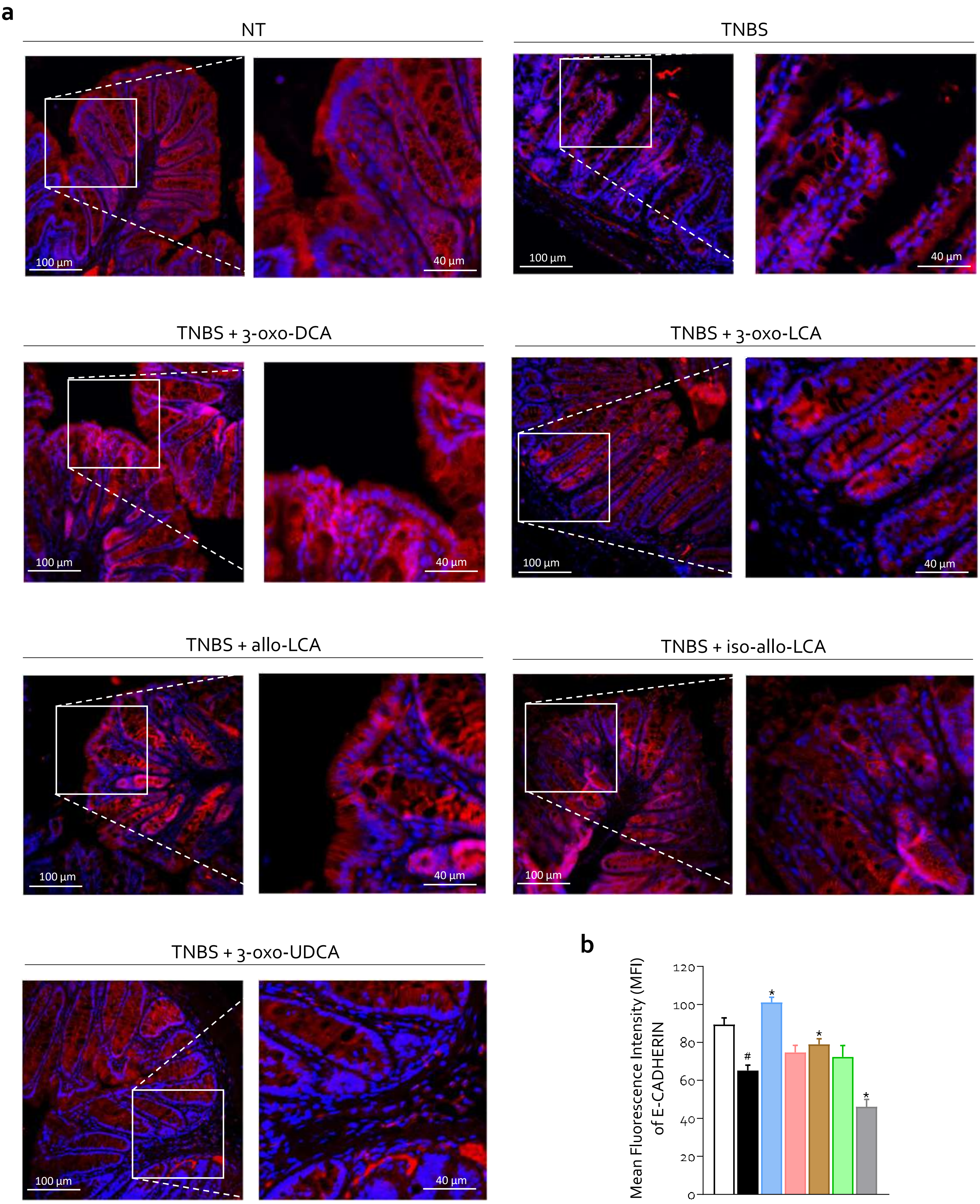
a
NT
TNBS
TNBS + 3-oxo-DCA
TNBS + 3-oxo-LCA
TNBS + allo-LCA
TNBS + iso-allo-LCA
TNBS + 3-oxo-UDCA
100 µm
40 µm
b
Mean Fluorescence Intensity (MFI) of E-CADHERIN
0
20
40
60
80
100
120
#
*

**Supplementary figure 2.** C57BL/6 male mice were treated with TNBS and oxo- or allo- bile acids derivates: 3-oxo-DCA, 3-oxo-LCA, allo -LCA, iso-allo-LCA, 3-oxo-UDCA (10 mg/Kg/daily). Evaluation of intestinal permeability alterations: (**a**) Immunofluorecent analysis of e-cadherin expression in colon sample (Magnification 40x and 100x). (**b**) Quantification of mean fluorescence intensity of e-cadherin. Data shown are the mean ± SEM of 3-7 mice per group. Statistical significance was assessed by 1-way ANOVA *$p \leq 0.05$. Abbreviations: NT, not treated; TNBS, 2,4,6-trinitrobenzenesulfonic acid; LCA, lithocholic acid; DCA, deoxycholic acid; UDCA, ursodeoxycholic acid.

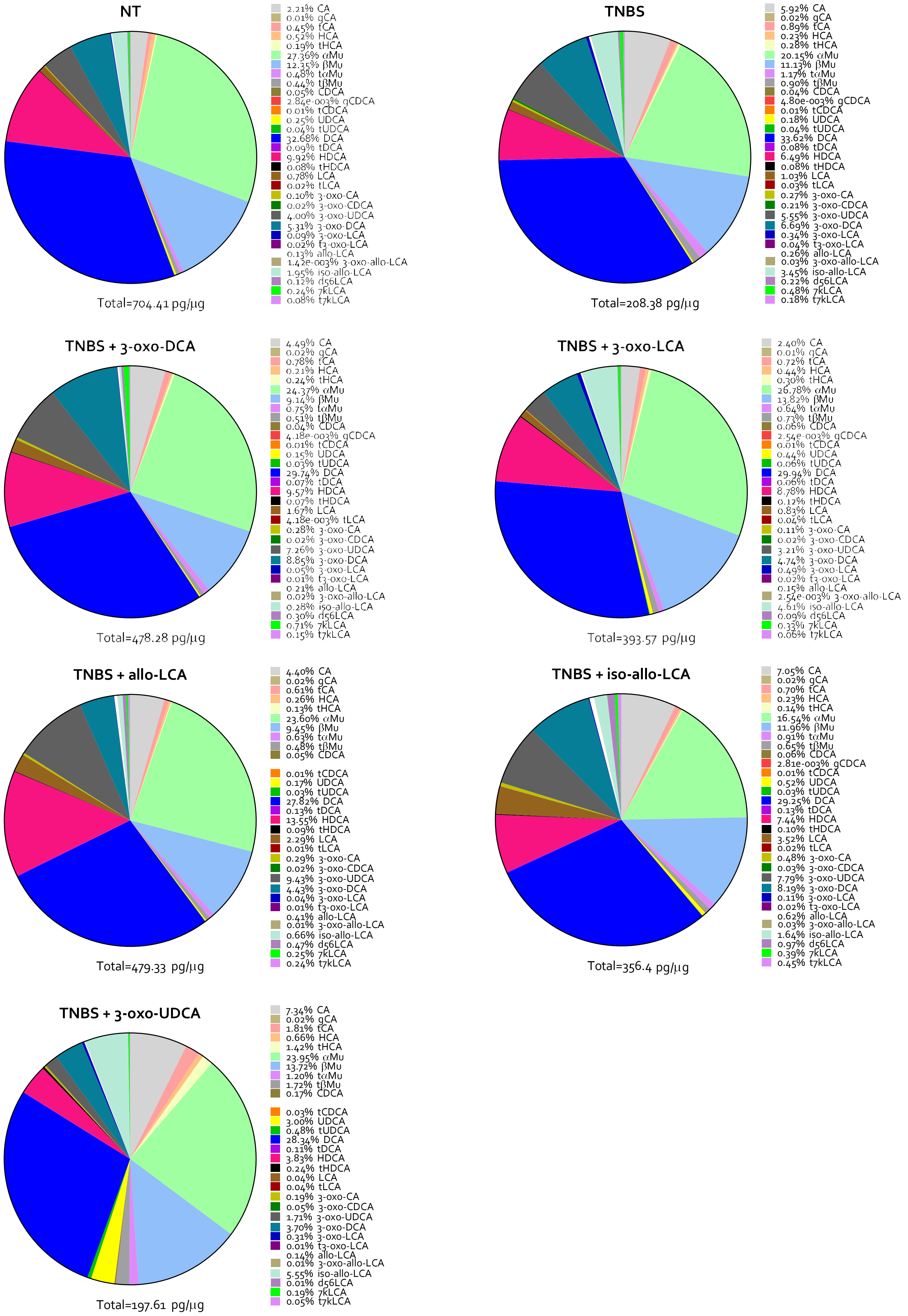
NT
2.21% CA
0.01% gCA
0.45% tCA
0.52% HCA
0.19% tHCA
27.36% αMu
12.35% βMu
0.48% tαMu
0.44% tβMu
0.05% CDCA
2.84e-003% gCDCA
0.01% tCDCA
0.25% UDCA
0.04% tUDCA
32.68% DCA
0.09% tDCA
9.92% HDCA
0.08% tHDCA
0.78% LCA
0.02% tLCA
0.10% 3-oxo-CA
0.02% 3-oxo-CDCA
4.00% 3-oxo-UDCA
5.31% 3-oxo-DCA
0.09% 3-oxo-LCA
0.02% t3-oxo-LCA
0.13% allo-LCA
1.42e-003% 3-oxo-allo-LCA
1.95% iso-allo-LCA
0.12% d56LCA
0.24% 7kLCA
0.08% t7kLCA
Total=704.41 pg/μg
TNBS
5.92% CA
0.02% gCA
0.89% tCA
0.23% HCA
0.28% tHCA
20.15% αMu
11.13% βMu
1.17% tαMu
0.90% tβMu
0.04% CDCA
4.80e-003% gCDCA
0.01% tCDCA
0.18% UDCA
0.04% tUDCA
33.62% DCA
0.08% tDCA
6.49% HDCA
0.08% tHDCA
1.03% LCA
0.03% tLCA
0.27% 3-oxo-CA
0.21% 3-oxo-CDCA
5.55% 3-oxo-UDCA
6.69% 3-oxo-DCA
0.34% 3-oxo-LCA
0.04% t3-oxo-LCA
0.26% allo-LCA
0.03% 3-oxo-allo-LCA
3.45% iso-allo-LCA
0.22% d56LCA
0.48% 7kLCA
0.18% t7kLCA
Total=208.38 pg/μg
TNBS + 3-oxo-DCA
4.49% CA
0.02% gCA
0.78% tCA
0.21% HCA
0.24% tHCA
24.37% αMu
9.14% βMu
0.75% tαMu
0.51% tβMu
0.04% CDCA
4.18e-003% gCDCA
0.01% tCDCA
0.15% UDCA
0.03% tUDCA
29.74% DCA
0.07% tDCA
9.57% HDCA
0.07% tHDCA
1.67% LCA
4.18e-003% tLCA
0.28% 3-oxo-CA
0.02% 3-oxo-CDCA
7.26% 3-oxo-UDCA
8.85% 3-oxo-DCA
0.05% 3-oxo-LCA
0.01% t3-oxo-LCA
0.21% allo-LCA
0.02% 3-oxo-allo-LCA
0.28% iso-allo-LCA
0.30% d56LCA
0.71% 7kLCA
0.15% t7kLCA
Total=478.28 pg/μg
TNBS + 3-oxo-LCA
2.40% CA
0.01% gCA
0.72% tCA
0.44% HCA
0.30% tHCA
26.78% αMu
13.82% βMu
0.64% tαMu
0.73% tβMu
0.06% CDCA
2.54e-003% gCDCA
0.01% tCDCA
0.44% UDCA
0.06% tUDCA
29.94% DCA
0.06% tDCA
8.78% HDCA
0.12% tHDCA
0.83% LCA
0.04% tLCA
0.11% 3-oxo-CA
0.02% 3-oxo-CDCA
3.21% 3-oxo-UDCA
4.74% 3-oxo-DCA
0.49% 3-oxo-LCA
0.02% t3-oxo-LCA
0.15% allo-LCA
2.54e-003% 3-oxo-allo-LCA
4.61% iso-allo-LCA
0.09% d56LCA
0.33% 7kLCA
0.06% t7kLCA
Total=393.57 pg/μg
TNBS + allo-LCA
4.40% CA
0.02% gCA
0.61% tCA
0.26% HCA
0.13% tHCA
23.60% αMu
9.45% βMu
0.63% tαMu
0.48% tβMu
0.05% CDCA
0.01% tCDCA
0.17% UDCA
0.03% tUDCA
27.82% DCA
0.13% tDCA
13.55% HDCA
0.09% tHDCA
2.29% LCA
0.01% tLCA
0.29% 3-oxo-CA
0.02% 3-oxo-CDCA
9.43% 3-oxo-UDCA
4.43% 3-oxo-DCA
0.04% 3-oxo-LCA
0.01% t3-oxo-LCA
0.41% allo-LCA
0.01% 3-oxo-allo-LCA
0.66% iso-allo-LCA
0.47% d56LCA
0.25% 7kLCA
0.24% t7kLCA
Total=479.33 pg/μg
TNBS + iso-allo-LCA
7.05% CA
0.02% gCA
0.70% tCA
0.23% HCA
0.14% tHCA
16.54% αMu
11.96% βMu
0.91% tαMu
0.65% tβMu
0.06% CDCA
2.81e-003% gCDCA
0.01% tCDCA
0.52% UDCA
0.03% tUDCA
29.25% DCA
0.13% tDCA
7.44% HDCA
0.10% tHDCA
3.52% LCA
0.02% tLCA
0.48% 3-oxo-CA
0.03% 3-oxo-CDCA
7.79% 3-oxo-UDCA
8.19% 3-oxo-DCA
0.11% 3-oxo-LCA
0.02% t3-oxo-LCA
0.62% allo-LCA
0.03% 3-oxo-allo-LCA
1.64% iso-allo-LCA
0.97% d56LCA
0.39% 7kLCA
0.45% t7kLCA
Total=356.4 pg/μg
TNBS + 3-oxo-UDCA
7.34% CA
0.02% gCA
1.81% tCA
0.66% HCA
1.42% tHCA
23.95% αMu
13.72% βMu
1.20% tαMu
1.72% tβMu
0.17% CDCA
0.03% tCDCA
3.00% UDCA
0.48% tUDCA
28.34% DCA
0.11% tDCA
3.83% HDCA
0.24% tHDCA
0.04% LCA
0.04% tLCA
0.19% 3-oxo-CA
0.05% 3-oxo-CDCA
1.71% 3-oxo-UDCA
3.70% 3-oxo-DCA
0.31% 3-oxo-LCA
0.01% t3-oxo-LCA
0.14% allo-LCA
0.01% 3-oxo-allo-LCA
5.55% iso-allo-LCA
0.01% d56LCA
0.19% 7kLCA
0.05% t7kLCA
Total=197.61 pg/μg

**Supplementary figure 3.** C57BL/6 male mice were treated with TNBS and oxo- or allo- bile acids derivates: 3-oxo-DCA, 3-oxo-LCA, allo-LCA, iso-allo-LCA, 3-oxo-UDCA (10 mg/Kg/daily). Histogram of fecal content of total bile acids evaluated in each experimental group. Abbreviations: NT, not treated; TNBS, 2,4,6-trinitrobenzenesulfonic acid; T, tauro; G, glyco; LCA, lithocholic acid; DCA, deoxycholic acid; CDCA, chenodeoxycholic acid; HDCA, hyodeoxycholic acid; UDCA, ursodeoxycholic acid; HCA, hyocholic acid; αMu, alpha-muricholic acid; βMu, beta-muricholic acid; 7k, 7keto; CA, cholic acid.

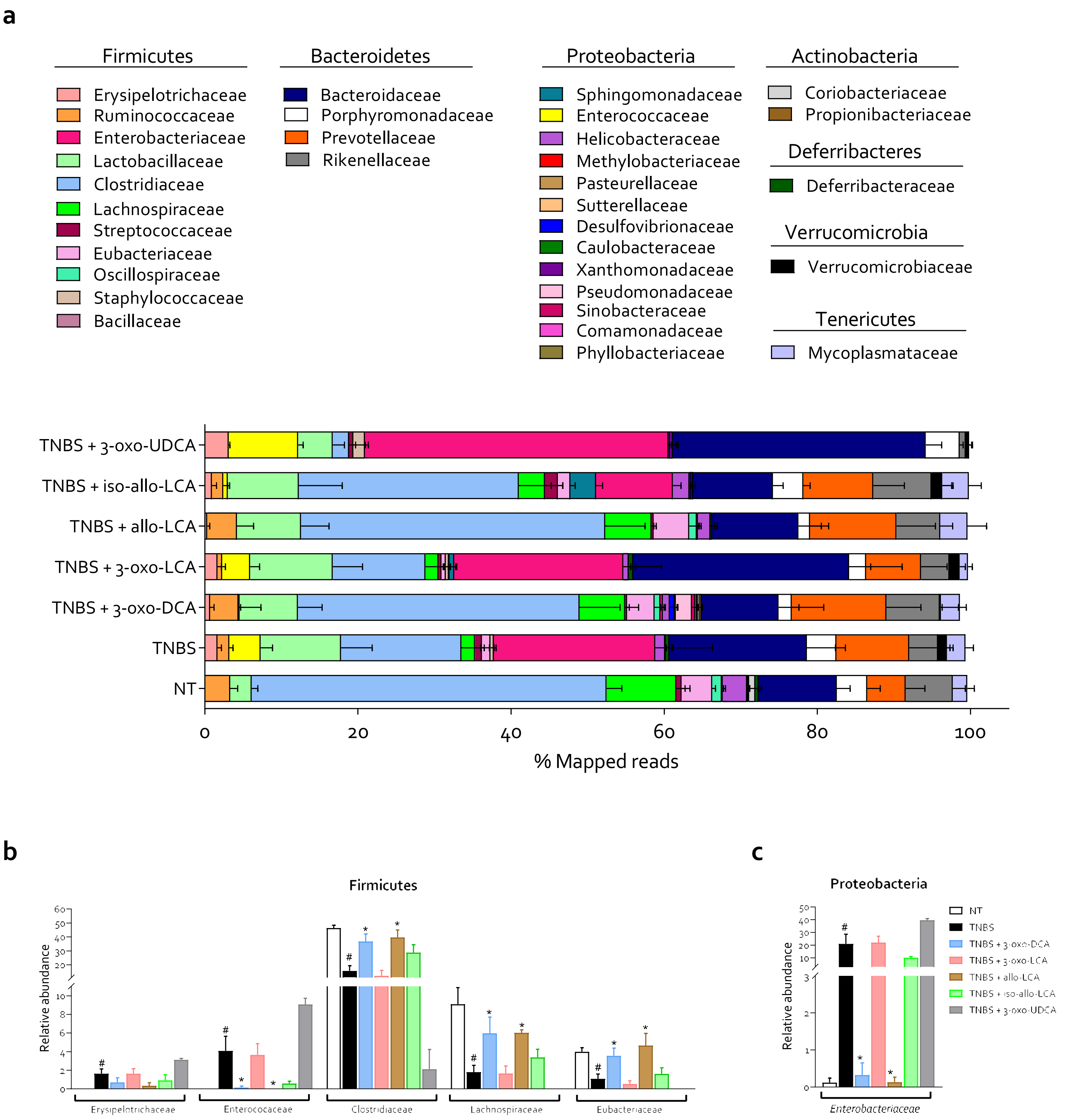
a
Firmicutes
Erysipelotrichaceae
Ruminococcaceae
Enterobacteriaceae
Lactobacillaceae
Clostridiaceae
Lachnospiraceae
Streptococcaceae
Eubacteriaceae
Oscillospiraceae
Staphylococcaceae
Bacillaceae
Bacteroidetes
Bacteroidaceae
Porphyromonadaceae
Prevotellaceae
Rikenellaceae
Proteobacteria
Sphingomonadaceae
Enterococcaceae
Helicobacteraceae
Methylobacteriaceae
Pasteurellaceae
Sutterellaceae
Desulfovibrionaceae
Caulobacteraceae
Xanthomonadaceae
Pseudomonadaceae
Sinobacteraceae
Comamonadaceae
Phyllobacteriaceae
Actinobacteria
Coriobacteriaceae
Propionibacteriaceae
Deferribacteres
Deferribacteraceae
Verrucomicrobia
Verrucomicrobiaceae
Tenericutes
Mycoplasmataceae
TNBS + 3-oxo-UDCA
TNBS + iso-allo-LCA
TNBS + allo-LCA
TNBS + 3-oxo-LCA
TNBS + 3-oxo-DCA
TNBS
NT
0
20
40
60
80
100
% Mapped reads
b
Firmicutes
Relative abundance
Erysipelotrichaceae
Enterococcaceae
Clostridiaceae
Lachnospiraceae
Eubacteriaceae
c
Proteobacteria
Relative abundance
Enterobacteriaceae
NT
TNBS
TNBS + 3-oxo-DCA
TNBS + 3-oxo-LCA
TNBS + allo-LCA
TNBS + iso-allo-LCA
TNBS + 3-oxo-UDCA

**Supplementary figure 4.** C57BL/6 male mice were treated with TNBS and oxo- or allo- bile acids derivates: 3-oxo-DCA, 3-oxo-LCA, allo-LCA, iso-allo-LCA, 3-oxo-UDCA (10 mg/Kg/daily). Analysis of fecal microbiota composition: (**a**) Relative abundance of all bacterial families evaluated each experimental group and calculated as percent of Mapped Reads; (**b,c**) relative abundance of families statistically modulated by TNBS. Data shown are the mean ± SEM of 3-7 mice per group. Statistical significance was assessed by 1-way ANOVA $*p \leq 0.05$. Abbreviations: NT, not treated; TNBS, 2,4,6-trinitrobenzenesulfonic acid; LCA, lithocholic acid; DCA, deoxycholic acid; UDCA, ursodeoxycholic acid.

| Phylum | Species | Relative abundance | | | | | | | | | | | | | |
|---|---|---|---|---|---|---|---|---|---|---|---|---|---|---|---|
| | | NT | | TNBS | | TNBS + 3-oxo-DCA | | TNBS + 3-oxo-LCA | | TNBS + Allo-LCA | | TNBS + Iso-Allo-LCA | | TNBS + 3-oxo-UDCA | |
| | | Mean | SEM | Mean | SEM | Mean | SEM | Mean | SEM | Mean | SEM | Mean | SEM | Mean | SEM |
| Actinobacteria | *Adlercreutzia equolifaciens* | 0,099 | 0,099 | 0,000 | 0,000 | 0,000 | 0,000 | 0,000 | 0,000 | 0,000 | 0,000 | 0,000 | 0,000 | 0,000 | 0,000 |
| Actinobacteria | *Enterorhabdus caecimuris* | 0,720 | 0,468 | 0,000 | 0,000 | 0,244 | 0,244 | 0,000 | 0,000 | 0,000 | 0,000 | 0,000 | 0,000 | 0,000 | 0,000 |
| Actinobacteria | *Propionibacterium acnes* | 0,041 | 0,041 | 0,124 | 0,087 | 0,294 | 0,126 | 0,000 | 0,000 | 0,000 | 0,000 | 0,000 | 0,000 | 0,000 | 0,000 |
| Bacteroidetes | *Alistipes putredinis* | 2,479 | 0,832 | 1,520 | 0,443 | 3,024 | 1,268 | 1,370 | 0,452 | 2,033 | 0,703 | 3,888 | 1,739 | 0,217 | 0,217 |
| Bacteroidetes | *Alistipes onderdonkii* | 0,369 | 0,146 | 0,190 | 0,074 | 0,256 | 0,159 | 0,364 | 0,164 | 0,345 | 0,125 | 0,553 | 0,252 | 0,000 | 0,000 |
| Bacteroidetes | *Alistipes finegoldii* | 1,474 | 0,324 | 1,099 | 0,225 | 1,510 | 0,587 | 1,061 | 0,192 | 1,948 | 0,462 | 1,585 | 0,393 | 0,380 | 0,380 |
| Bacteroidetes | *Alistipes sp,* | 0,194 | 0,096 | 0,154 | 0,075 | 0,222 | 0,142 | 0,339 | 0,025 | 0,343 | 0,120 | 0,333 | 0,113 | 0,117 | 0,117 |
| Bacteroidetes | *Alistipes massiliensis* | 1,010 | 0,215 | 0,529 | 0,247 | 1,540 | 0,552 | 0,354 | 0,172 | 1,003 | 0,357 | 1,075 | 0,395 | 0,000 | 0,000 |
| Bacteroidetes | *Alistipes senegalensis* | 0,000 | 0,000 | 0,054 | 0,054 | 0,076 | 0,076 | 0,049 | 0,049 | 0,000 | 0,000 | 0,153 | 0,153 | 0,000 | 0,000 |
| Bacteroidetes | *Bacteroides sp,* | 3,261 | 0,673 | 3,119 | 0,525 | 3,004 | 1,552 | 4,859 | 0,899 | 4,688 | 1,890 | 2,735 | 0,434 | 6,607 | 1,560 |
| Bacteroidetes | *Bacteroides ovatus* | 0,364 | 0,096 | 1,149 | 0,462 | 0,460 | 0,460 | 2,011 | 0,606 | 0,363 | 0,148 | 0,423 | 0,255 | 1,797 | 0,258 |
| Bacteroidetes | *Bacteroides thetaiotaomicron* | 0,000 | 0,000 | 1,740 | 1,033 | 0,102 | 0,102 | 1,863 | 0,668 | 0,120 | 0,120 | 0,698 | 0,403 | 3,280 | 0,530 |
| Bacteroidetes | *Bacteroides acidifaciens* | 5,631 | 1,162 | 10,754 | 2,761 | 6,040 | 3,849 | 17,951 | 5,272 | 5,715 | 2,430 | 6,258 | 0,813 | 19,707 | 2,845 |
| Bacteroidetes | *Bacteroides uniformis* | 0,137 | 0,090 | 0,833 | 0,430 | 0,060 | 0,060 | 0,591 | 0,542 | 0,303 | 0,303 | 0,125 | 0,125 | 0,243 | 0,243 |
| Bacteroidetes | *Bacteroides caccae* | 0,000 | 0,000 | 0,104 | 0,104 | 0,000 | 0,000 | 0,271 | 0,271 | 0,000 | 0,000 | 0,000 | 0,000 | 1,253 | 0,683 |
| Bacteroidetes | *Bacteroides salyersiae* | 0,000 | 0,000 | 0,000 | 0,000 | 0,000 | 0,000 | 0,429 | 0,429 | 0,000 | 0,000 | 0,000 | 0,000 | 0,000 | 0,000 |
| Bacteroidetes | *Bacteroides stercoris* | 0,000 | 0,000 | 0,000 | 0,000 | 0,000 | 0,000 | 0,096 | 0,096 | 0,000 | 0,000 | 0,000 | 0,000 | 0,000 | 0,000 |
| Bacteroidetes | *Bacteroides nordii* | 0,000 | 0,000 | 0,000 | 0,000 | 0,000 | 0,000 | 0,091 | 0,091 | 0,000 | 0,000 | 0,000 | 0,000 | 0,000 | 0,000 |
| Bacteroidetes | *Bacteroides sartorii* | 0,650 | 0,530 | 0,000 | 0,000 | 0,284 | 0,284 | 0,000 | 0,000 | 0,000 | 0,000 | 0,218 | 0,218 | 0,000 | 0,000 |
| Bacteroidetes | *Bacteroides finegoldii* | 0,000 | 0,000 | 0,170 | 0,170 | 0,000 | 0,000 | 0,049 | 0,049 | 0,000 | 0,000 | 0,000 | 0,000 | 0,113 | 0,113 |
| Bacteroidetes | *Bacteroides chinchillae* | 0,000 | 0,000 | 0,000 | 0,000 | 0,050 | 0,050 | 0,000 | 0,000 | 0,000 | 0,000 | 0,000 | 0,000 | 0,000 | 0,000 |
| Bacteroidetes | *Bacteroides faecichinchillae* | 0,103 | 0,103 | 0,000 | 0,000 | 0,000 | 0,000 | 0,000 | 0,000 | 0,000 | 0,000 | 0,000 | 0,000 | 0,000 | 0,000 |
| Bacteroidetes | *Barnesiella intestinihominis* | 0,000 | 0,000 | 0,344 | 0,344 | 0,000 | 0,000 | 0,000 | 0,000 | 1,108 | 1,108 | 0,000 | 0,000 | 0,000 | 0,000 |
| Bacteroidetes | *Odoribacter splanchnicus* | 2,043 | 1,596 | 0,189 | 0,124 | 1,036 | 1,036 | 0,651 | 0,427 | 0,000 | 0,000 | 1,003 | 0,664 | 0,000 | 0,000 |
| Bacteroidetes | *Odoribacter laneus* | 0,477 | 0,477 | 0,110 | 0,110 | 0,000 | 0,000 | 0,000 | 0,000 | 0,000 | 0,000 | 0,000 | 0,000 | 0,000 | 0,000 |
| Bacteroidetes | *Parabacteroides johnsonii* | 0,197 | 0,197 | 0,356 | 0,172 | 0,000 | 0,000 | 0,126 | 0,083 | 0,148 | 0,148 | 0,663 | 0,392 | 0,000 | 0,000 |
| Bacteroidetes | *Parabacteroides merdae* | 0,743 | 0,452 | 1,088 | 0,370 | 0,180 | 0,180 | 0,416 | 0,276 | 0,248 | 0,248 | 2,123 | 0,492 | 0,000 | 0,000 |
| Bacteroidetes | *Parabacteroides distasonis* | 0,534 | 0,364 | 1,083 | 0,571 | 0,492 | 0,327 | 0,316 | 0,160 | 0,000 | 0,000 | 0,153 | 0,153 | 1,847 | 0,542 |
| Bacteroidetes | *Parabacteroides sp,* | 0,000 | 0,000 | 0,251 | 0,165 | 0,000 | 0,000 | 0,000 | 0,000 | 0,000 | 0,000 | 0,000 | 0,000 | 0,217 | 0,217 |
| Bacteroidetes | *Parabacteroides goldsteinii* | 0,000 | 0,000 | 0,420 | 0,246 | 0,000 | 0,000 | 0,714 | 0,418 | 0,000 | 0,000 | 0,000 | 0,000 | 2,237 | 0,388 |
| Bacteroidetes | *Paraprevotella xylaniphila* | 1,021 | 1,021 | 1,034 | 0,678 | 0,000 | 0,000 | 0,000 | 0,000 | 0,860 | 0,860 | 0,000 | 0,000 | 0,000 | 0,000 |
| Bacteroidetes | *Prevotella loescheii* | 0,000 | 0,000 | 1,261 | 0,934 | 0,252 | 0,252 | 0,320 | 0,320 | 0,333 | 0,333 | 0,903 | 0,903 | 0,000 | 0,000 |
| Bacteroidetes | *Prevotella sp,* | 3,151 | 2,054 | 3,944 | 2,560 | 8,236 | 4,382 | 4,600 | 2,314 | 7,455 | 5,809 | 4,740 | 2,570 | 0,000 | 0,000 |
| Bacteroidetes | *Prevotella genomosp,* | 0,186 | 0,186 | 1,204 | 0,811 | 0,350 | 0,230 | 0,311 | 0,202 | 1,058 | 0,688 | 1,325 | 1,325 | 0,000 | 0,000 |
| Bacteroidetes | *Prevotella Prevotella* | 0,653 | 0,653 | 1,888 | 1,888 | 3,282 | 1,648 | 1,946 | 1,307 | 1,605 | 1,605 | 2,175 | 2,175 | 0,000 | 0,000 |
| Bacteroidetes | *Prevotella shahii* | 0,000 | 0,000 | 0,118 | 0,118 | 0,118 | 0,072 | 0,000 | 0,000 | 0,000 | 0,000 | 0,000 | 0,000 | 0,000 | 0,000 |
| Bacteroidetes | *Prevotella buccae* | 0,000 | 0,000 | 0,066 | 0,066 | 0,132 | 0,132 | 0,000 | 0,000 | 0,000 | 0,000 | 0,000 | 0,000 | 0,000 | 0,000 |
| Bacteroidetes | *Rikenella microfusus* | 0,416 | 0,416 | 0,088 | 0,088 | 0,216 | 0,216 | 0,150 | 0,150 | 0,000 | 0,000 | 0,000 | 0,000 | 0,000 | 0,000 |
| Deferribacteres | *Mucispirillum schaedleri* | 0,503 | 0,341 | 0,588 | 0,459 | 0,056 | 0,056 | 0,569 | 0,324 | 0,248 | 0,248 | 0,000 | 0,000 | 0,100 | 0,100 |
| Firmicutes | *Anaerostipes hadrus* | 0,583 | 0,204 | 0,000 | 0,000 | 0,000 | 0,000 | 0,000 | 0,000 | 0,000 | 0,000 | 0,000 | 0,000 | 0,000 | 0,000 |
| Firmicutes | *Anaerostipes sp,* | 0,237 | 0,117 | 0,000 | 0,000 | 0,000 | 0,000 | 0,000 | 0,000 | 0,000 | 0,000 | 0,000 | 0,000 | 0,000 | 0,000 |
| Firmicutes | *Anaerotruncus sp,* | 0,201 | 0,201 | 0,000 | 0,000 | 0,000 | 0,000 | 0,000 | 0,000 | 0,000 | 0,000 | 0,000 | 0,000 | 0,000 | 0,000 |
| Firmicutes | *Anaerotruncus colihominis* | 0,563 | 0,563 | 0,000 | 0,000 | 0,300 | 0,300 | 0,000 | 0,000 | 0,000 | 0,000 | 0,000 | 0,000 | 0,000 | 0,000 |
| Firmicutes | *Bacillus sp,* | 0,000 | 0,000 | 0,000 | 0,000 | 0,000 | 0,000 | 0,046 | 0,046 | 0,000 | 0,000 | 0,000 | 0,000 | 0,000 | 0,000 |

| | | | | | | | | | | | | | | | |
|---|---|---|---|---|---|---|---|---|---|---|---|---|---|---|---|
| Firmicutes | *Blautia gnavus* | 1,601 | 0,225 | 0,850 | 0,354 | 1,960 | 0,336 | 0,910 | 0,367 | 1,983 | 0,258 | 0,470 | 0,470 | 0,000 | 0,000 |
| Firmicutes | *Blautia torques* | 0,081 | 0,081 | 0,000 | 0,000 | 0,284 | 0,284 | 0,000 | 0,000 | 0,213 | 0,213 | 0,000 | 0,000 | 0,000 | 0,000 |
| Firmicutes | *Blautia coccoides* | 0,071 | 0,071 | 0,000 | 0,000 | 0,112 | 0,112 | 0,000 | 0,000 | 0,000 | 0,000 | 0,000 | 0,000 | 0,000 | 0,000 |
| Firmicutes | *Blautia sp,* | 0,990 | 0,525 | 0,000 | 0,000 | 0,240 | 0,240 | 0,254 | 0,254 | 0,318 | 0,318 | 0,000 | 0,000 | 0,000 | 0,000 |
| Firmicutes | *Blautia producta* | 0,039 | 0,039 | 0,000 | 0,000 | 0,000 | 0,000 | 0,000 | 0,000 | 0,000 | 0,000 | 0,000 | 0,000 | 0,000 | 0,000 |
| Firmicutes | *Blautia obeum* | 0,267 | 0,198 | 0,000 | 0,000 | 0,000 | 0,000 | 0,000 | 0,000 | 0,305 | 0,305 | 0,000 | 0,000 | 0,000 | 0,000 |
| Firmicutes | *Butyricicoccus pullicaecorum* | 0,914 | 0,615 | 0,000 | 0,000 | 0,000 | 0,000 | 0,000 | 0,000 | 0,925 | 0,925 | 0,000 | 0,000 | 0,000 | 0,000 |
| Firmicutes | *Candidatus Arthromitus* | 2,447 | 1,103 | 0,513 | 0,513 | 0,000 | 0,000 | 0,151 | 0,151 | 1,473 | 1,152 | 0,400 | 0,400 | 0,000 | 0,000 |
| Firmicutes | *Candidatus Soleaferrea* | 0,000 | 0,000 | 0,000 | 0,000 | 0,000 | 0,000 | 0,000 | 0,000 | 0,000 | 0,000 | 0,000 | 0,000 | 0,000 | 0,000 |
| Firmicutes | *Clostridium sp,* | 41,443 | 2,216 | 14,880 | 3,279 | 35,802 | 5,337 | 11,679 | 3,651 | 36,380 | 4,261 | 27,833 | 5,257 | 2,117 | 2,117 |
| Firmicutes | *Clostridium fusiformis* | 1,571 | 0,431 | 0,316 | 0,110 | 0,956 | 0,185 | 0,271 | 0,180 | 0,948 | 0,275 | 0,530 | 0,314 | 0,000 | 0,000 |
| Firmicutes | *Coprobacillus cateniformis* | 0,000 | 0,000 | 0,398 | 0,206 | 0,000 | 0,000 | 0,204 | 0,204 | 0,333 | 0,333 | 0,175 | 0,175 | 1,073 | 1,073 |
| Firmicutes | *Dorea formicigenerans* | 0,000 | 0,000 | 0,000 | 0,000 | 0,058 | 0,058 | 0,000 | 0,000 | 0,000 | 0,000 | 0,000 | 0,000 | 0,000 | 0,000 |
| Firmicutes | *Enterococcus faecalis* | 0,000 | 0,000 | 2,208 | 0,747 | 0,134 | 0,134 | 1,823 | 0,480 | 0,000 | 0,000 | 0,573 | 0,243 | 5,070 | 0,867 |
| Firmicutes | *Enterococcus sp,* | 0,000 | 0,000 | 0,180 | 0,088 | 0,000 | 0,000 | 0,171 | 0,089 | 0,000 | 0,000 | 0,000 | 0,000 | 0,470 | 0,104 |
| Firmicutes | *Enterococcus faecium* | 0,000 | 0,000 | 0,480 | 0,241 | 0,000 | 0,000 | 0,316 | 0,132 | 0,000 | 0,000 | 0,000 | 0,000 | 0,850 | 0,270 |
| Firmicutes | *Enterococcus gallinarum* | 0,000 | 0,000 | 0,980 | 0,420 | 0,000 | 0,000 | 1,090 | 0,420 | 0,000 | 0,000 | 0,000 | 0,000 | 2,030 | 0,825 |
| Firmicutes | *Enterococcus casseliflavus* | 0,000 | 0,000 | 0,173 | 0,088 | 0,000 | 0,000 | 0,129 | 0,083 | 0,000 | 0,000 | 0,000 | 0,000 | 0,433 | 0,433 |
| Firmicutes | *Enterococcus durans* | 0,000 | 0,000 | 0,000 | 0,000 | 0,000 | 0,000 | 0,043 | 0,043 | 0,000 | 0,000 | 0,000 | 0,000 | 0,000 | 0,000 |
| Firmicutes | *Erysipelatoclostridium cocleatum* | 0,000 | 0,000 | 1,245 | 0,481 | 0,688 | 0,510 | 1,437 | 0,444 | 0,000 | 0,000 | 0,745 | 0,610 | 2,050 | 1,034 |
| Firmicutes | *Eubacterium plexicaudatum* | 2,154 | 0,383 | 0,404 | 0,201 | 1,940 | 0,306 | 0,389 | 0,254 | 2,613 | 0,797 | 1,358 | 0,600 | 0,000 | 0,000 |
| Firmicutes | *Eubacterium ramulus* | 1,046 | 0,259 | 0,349 | 0,233 | 0,918 | 0,549 | 0,143 | 0,143 | 0,963 | 0,703 | 0,260 | 0,260 | 0,000 | 0,000 |
| Firmicutes | *Eubacterium coprostanoligenes* | 0,387 | 0,262 | 0,344 | 0,261 | 0,594 | 0,368 | 0,000 | 0,000 | 1,070 | 0,847 | 0,000 | 0,000 | 0,000 | 0,000 |
| Firmicutes | *Eubacterium oxidoreducens* | 0,400 | 0,165 | 0,000 | 0,000 | 0,110 | 0,110 | 0,000 | 0,000 | 0,000 | 0,000 | 0,000 | 0,000 | 0,000 | 0,000 |
| Firmicutes | *Lachnoclostridium hathewayi* | 0,391 | 0,285 | 0,000 | 0,000 | 0,206 | 0,206 | 0,000 | 0,000 | 0,493 | 0,323 | 0,000 | 0,000 | 0,000 | 0,000 |
| Firmicutes | *Lachnoclostridium celerecrescens* | 0,290 | 0,156 | 0,000 | 0,000 | 0,106 | 0,106 | 0,000 | 0,000 | 0,128 | 0,128 | 0,000 | 0,000 | 0,000 | 0,000 |
| Firmicutes | *Lachnoclostridium clostridioforme* | 0,836 | 0,414 | 0,000 | 0,000 | 0,000 | 0,000 | 0,000 | 0,000 | 0,000 | 0,000 | 0,000 | 0,000 | 0,000 | 0,000 |
| Firmicutes | *Lachnoclostridium scindens* | 0,556 | 0,374 | 0,099 | 0,099 | 0,120 | 0,120 | 0,000 | 0,000 | 0,000 | 0,000 | 0,415 | 0,245 | 0,000 | 0,000 |
| Firmicutes | *Lachnoclostridium fissicatena* | 0,059 | 0,059 | 0,000 | 0,000 | 0,240 | 0,147 | 0,090 | 0,090 | 0,075 | 0,075 | 0,000 | 0,000 | 0,000 | 0,000 |
| Firmicutes | *Lachnoclostridium hylemonae* | 0,301 | 0,203 | 0,000 | 0,000 | 0,000 | 0,000 | 0,000 | 0,000 | 0,000 | 0,000 | 0,000 | 0,000 | 0,000 | 0,000 |
| Firmicutes | *Lachnoclostridium symbiosum* | 0,000 | 0,000 | 0,000 | 0,000 | 0,000 | 0,000 | 0,000 | 0,000 | 0,170 | 0,170 | 0,000 | 0,000 | 0,000 | 0,000 |
| Firmicutes | *Lachnoclostridium citroniae* | 0,000 | 0,000 | 0,000 | 0,000 | 0,000 | 0,000 | 0,000 | 0,000 | 0,000 | 0,000 | 0,000 | 0,000 | 0,000 | 0,000 |
| Firmicutes | *Lachnospira pectinoschiza* | 0,426 | 0,333 | 0,000 | 0,000 | 0,204 | 0,204 | 0,000 | 0,000 | 0,000 | 0,000 | 0,000 | 0,000 | 0,000 | 0,000 |
| Firmicutes | *Lactobacillus animalis* | 0,389 | 0,132 | 2,124 | 0,871 | 1,664 | 0,694 | 1,553 | 0,647 | 1,648 | 0,826 | 1,955 | 1,278 | 0,877 | 0,292 |
| Firmicutes | *Lactobacillus murinus* | 1,530 | 0,385 | 8,064 | 3,149 | 5,956 | 2,448 | 5,761 | 2,138 | 6,355 | 2,531 | 7,298 | 4,434 | 3,663 | 1,211 |
| Firmicutes | *Lactobacillus gasseri* | 0,140 | 0,095 | 0,040 | 0,040 | 0,000 | 0,000 | 0,690 | 0,690 | 0,000 | 0,000 | 0,000 | 0,000 | 0,000 | 0,000 |
| Firmicutes | *Lactobacillus reuteri* | 0,156 | 0,156 | 0,000 | 0,000 | 0,000 | 0,000 | 0,980 | 0,980 | 0,000 | 0,000 | 0,000 | 0,000 | 0,000 | 0,000 |
| Firmicutes | *Lactobacillus johnsonii* | 0,513 | 0,282 | 0,120 | 0,120 | 0,000 | 0,000 | 1,599 | 1,599 | 0,000 | 0,000 | 0,000 | 0,000 | 0,000 | 0,000 |
| Firmicutes | *Lactobacillus apodemi* | 0,000 | 0,000 | 0,178 | 0,178 | 0,000 | 0,000 | 0,184 | 0,184 | 0,328 | 0,328 | 0,000 | 0,000 | 0,000 | 0,000 |
| Firmicutes | *Lactococcus lactis* | 0,000 | 0,000 | 0,596 | 0,269 | 0,414 | 0,241 | 0,377 | 0,204 | 0,310 | 0,310 | 0,545 | 0,394 | 0,193 | 0,193 |
| Firmicutes | *Lactococcus garvieae* | 0,627 | 0,418 | 0,000 | 0,000 | 0,000 | 0,000 | 0,000 | 0,000 | 0,000 | 0,000 | 0,648 | 0,648 | 0,000 | 0,000 |
| Firmicutes | *Lactococcus sp,* | 0,000 | 0,000 | 0,000 | 0,000 | 0,000 | 0,000 | 0,000 | 0,000 | 0,000 | 0,000 | 0,075 | 0,075 | 0,000 | 0,000 |
| Firmicutes | *Lysinibacillus fusiformis* | 0,000 | 0,000 | 0,000 | 0,000 | 0,000 | 0,000 | 0,043 | 0,043 | 0,000 | 0,000 | 0,000 | 0,000 | 0,000 | 0,000 |
| Firmicutes | *Marvinbryantia formatexigens* | 1,581 | 0,349 | 0,696 | 0,280 | 1,662 | 0,632 | 0,411 | 0,269 | 1,908 | 0,535 | 1,205 | 0,402 | 0,000 | 0,000 |
| Firmicutes | *Oscillibacter sp,* | 0,801 | 0,263 | 0,000 | 0,000 | 0,770 | 0,479 | 0,000 | 0,000 | 0,978 | 0,571 | 0,000 | 0,000 | 0,000 | 0,000 |
| Firmicutes | *Oscillibacter ruminantium* | 0,153 | 0,153 | 0,000 | 0,000 | 0,000 | 0,000 | 0,000 | 0,000 | 0,000 | 0,000 | 0,000 | 0,000 | 0,000 | 0,000 |
| Firmicutes | *Oscillibacter valericigenes* | 0,306 | 0,306 | 0,000 | 0,000 | 0,000 | 0,000 | 0,000 | 0,000 | 0,000 | 0,000 | 0,000 | 0,000 | 0,000 | 0,000 |

| | | | | | | | | | | | | | | | |
|---|---|---|---|---|---|---|---|---|---|---|---|---|---|---|---|
| Firmicutes | *Oscillibacter guilliermondii* | 0,614 | 0,614 | 0,000 | 0,000 | 1,348 | 1,348 | 0,000 | 0,000 | 1,850 | 1,850 | 0,000 | 0,000 | 0,000 | 0,000 |
| Firmicutes | *Papillibacter cinnamivorans* | 0,153 | 0,153 | 0,000 | 0,000 | 0,000 | 0,000 | 0,000 | 0,000 | 0,000 | 0,000 | 0,000 | 0,000 | 0,000 | 0,000 |
| Firmicutes | *Robinsoniella peoriensis* | 0,401 | 0,401 | 0,000 | 0,000 | 0,406 | 0,406 | 0,000 | 0,000 | 0,433 | 0,433 | 1,070 | 0,623 | 0,000 | 0,000 |
| Firmicutes | *Roseburia intestinalis* | 0,144 | 0,144 | 0,143 | 0,143 | 0,086 | 0,086 | 0,000 | 0,000 | 0,000 | 0,000 | 0,238 | 0,238 | 0,000 | 0,000 |
| Firmicutes | *Roseburia sp,* | 0,069 | 0,069 | 0,000 | 0,000 | 0,000 | 0,000 | 0,000 | 0,000 | 0,000 | 0,000 | 0,000 | 0,000 | 0,000 | 0,000 |
| Firmicutes | *Roseburia faecis* | 0,149 | 0,149 | 0,000 | 0,000 | 0,288 | 0,288 | 0,000 | 0,000 | 0,000 | 0,000 | 0,000 | 0,000 | 0,000 | 0,000 |
| Firmicutes | *Ruminiclostridium siraeum* | 0,363 | 0,182 | 0,118 | 0,118 | 0,296 | 0,296 | 0,000 | 0,000 | 0,095 | 0,095 | 0,000 | 0,000 | 0,000 | 0,000 |
| Firmicutes | *Ruminiclostridium viride* | 0,181 | 0,181 | 0,000 | 0,000 | 0,000 | 0,000 | 0,000 | 0,000 | 0,000 | 0,000 | 0,000 | 0,000 | 0,000 | 0,000 |
| Firmicutes | *Ruminiclostridium methylpentosum* | 0,000 | 0,000 | 0,000 | 0,000 | 0,536 | 0,536 | 0,000 | 0,000 | 0,000 | 0,000 | 0,000 | 0,000 | 0,000 | 0,000 |
| Firmicutes | *Ruminococcus flavefaciens* | 0,391 | 0,391 | 1,230 | 0,551 | 0,366 | 0,366 | 0,630 | 0,422 | 0,900 | 0,582 | 1,140 | 0,672 | 0,000 | 0,000 |
| Firmicutes | *Ruminococcus sp,* | 0,650 | 0,570 | 0,210 | 0,210 | 0,880 | 0,880 | 0,000 | 0,000 | 1,015 | 0,602 | 0,353 | 0,353 | 0,000 | 0,000 |
| Firmicutes | *Ruminococcus albus* | 0,210 | 0,210 | 0,000 | 0,000 | 0,000 | 0,000 | 0,000 | 0,000 | 0,000 | 0,000 | 0,000 | 0,000 | 0,000 | 0,000 |
| Firmicutes | *Staphylococcus sciuri* | 0,000 | 0,000 | 0,333 | 0,194 | 0,000 | 0,000 | 0,453 | 0,211 | 0,000 | 0,000 | 0,000 | 0,000 | 1,330 | 0,414 |
| Firmicutes | *Staphylococcus sp,* | 0,000 | 0,000 | 0,000 | 0,000 | 0,000 | 0,000 | 0,000 | 0,000 | 0,000 | 0,000 | 0,000 | 0,000 | 0,087 | 0,087 |
| Firmicutes | *Staphylococcus epidermidis* | 0,000 | 0,000 | 0,084 | 0,084 | 0,000 | 0,000 | 0,000 | 0,000 | 0,000 | 0,000 | 0,000 | 0,000 | 0,000 | 0,000 |
| Firmicutes | *Streptococcus sp,* | 0,000 | 0,000 | 0,334 | 0,219 | 0,000 | 0,000 | 0,119 | 0,077 | 0,000 | 0,000 | 0,445 | 0,285 | 0,353 | 0,353 |
| Proteobacteria | *Bradyrhizobium sp,* | 0,000 | 0,000 | 0,000 | 0,000 | 0,086 | 0,086 | 0,000 | 0,000 | 0,000 | 0,000 | 0,000 | 0,000 | 0,000 | 0,000 |
| Proteobacteria | *Brevundimonas sp,* | 0,000 | 0,000 | 0,000 | 0,000 | 0,000 | 0,000 | 0,000 | 0,000 | 0,000 | 0,000 | 0,000 | 0,000 | 0,000 | 0,000 |
| Proteobacteria | *Caulobacter sp,* | 0,061 | 0,061 | 0,083 | 0,055 | 0,192 | 0,079 | 0,000 | 0,000 | 0,000 | 0,000 | 0,000 | 0,000 | 0,000 | 0,000 |
| Proteobacteria | *Caulobacter vibrioides* | 0,000 | 0,000 | 0,000 | 0,000 | 0,086 | 0,086 | 0,000 | 0,000 | 0,000 | 0,000 | 0,000 | 0,000 | 0,000 | 0,000 |
| Proteobacteria | *Cronobacter sakazakii* | 0,000 | 0,000 | 0,000 | 0,000 | 0,000 | 0,000 | 0,000 | 0,000 | 0,000 | 0,000 | 0,000 | 0,000 | 0,000 | 0,000 |
| Proteobacteria | *Desulfovibrio sp,* | 0,167 | 0,109 | 0,000 | 0,000 | 0,640 | 0,399 | 0,000 | 0,000 | 0,000 | 0,000 | 0,300 | 0,300 | 0,000 | 0,000 |
| Proteobacteria | *Enhydrobacter aerosaccus* | 0,000 | 0,000 | 0,000 | 0,000 | 0,096 | 0,096 | 0,000 | 0,000 | 0,000 | 0,000 | 0,000 | 0,000 | 0,000 | 0,000 |
| Proteobacteria | *Enterobacter sp,* | 0,000 | 0,000 | 0,000 | 0,000 | 0,000 | 0,000 | 0,061 | 0,061 | 0,000 | 0,000 | 0,000 | 0,000 | 0,000 | 0,000 |
| Proteobacteria | *Escherichia coli* | 0,269 | 0,117 | 18,330 | 6,415 | 0,470 | 0,291 | 17,927 | 4,817 | 0,343 | 0,119 | 3,788 | 1,030 | 34,837 | 1,318 |
| Proteobacteria | *Helicobacter hepaticus* | 2,860 | 1,254 | 1,214 | 0,801 | 1,024 | 0,400 | 0,773 | 0,263 | 1,583 | 0,631 | 2,145 | 0,185 | 0,200 | 0,200 |
| Proteobacteria | *Helicobacter bilis* | 0,076 | 0,076 | 0,055 | 0,055 | 0,000 | 0,000 | 0,000 | 0,000 | 0,000 | 0,000 | 0,000 | 0,000 | 0,000 | 0,000 |
| Proteobacteria | *Helicobacter canadensis* | 0,050 | 0,050 | 0,000 | 0,000 | 0,000 | 0,000 | 0,000 | 0,000 | 0,000 | 0,000 | 0,000 | 0,000 | 0,000 | 0,000 |
| Proteobacteria | *Helicobacter sp,* | 0,131 | 0,086 | 0,000 | 0,000 | 0,000 | 0,000 | 0,000 | 0,000 | 0,173 | 0,173 | 0,000 | 0,000 | 0,000 | 0,000 |
| Proteobacteria | *Klebsiella sp,* | 0,000 | 0,000 | 0,000 | 0,000 | 0,000 | 0,000 | 0,000 | 0,000 | 0,000 | 0,000 | 0,000 | 0,000 | 0,130 | 0,130 |
| Proteobacteria | *Mesorhizobium sp,* | 0,000 | 0,000 | 0,000 | 0,000 | 0,136 | 0,136 | 0,000 | 0,000 | 0,000 | 0,000 | 0,000 | 0,000 | 0,000 | 0,000 |
| Proteobacteria | *Methylobacterium sp,* | 0,000 | 0,000 | 0,000 | 0,000 | 0,000 | 0,000 | 0,000 | 0,000 | 0,000 | 0,000 | 0,175 | 0,175 | 0,000 | 0,000 |
| Proteobacteria | *Nevskia soli* | 0,000 | 0,000 | 0,000 | 0,000 | 0,444 | 0,444 | 0,000 | 0,000 | 0,000 | 0,000 | 0,000 | 0,000 | 0,000 | 0,000 |
| Proteobacteria | *Pantoea sp,* | 0,000 | 0,000 | 0,000 | 0,000 | 0,000 | 0,000 | 0,054 | 0,054 | 0,000 | 0,000 | 0,460 | 0,048 | 0,000 | 0,000 |
| Proteobacteria | *Pantoea agglomerans* | 0,000 | 0,000 | 0,000 | 0,000 | 0,000 | 0,000 | 1,333 | 0,735 | 0,000 | 0,000 | 5,440 | 0,176 | 0,000 | 0,000 |
| Proteobacteria | *Parasutterella excrementihominis* | 0,000 | 0,000 | 0,000 | 0,000 | 0,160 | 0,100 | 0,000 | 0,000 | 0,000 | 0,000 | 0,110 | 0,110 | 0,000 | 0,000 |
| Proteobacteria | *Pasteurella sp,* | 0,000 | 0,000 | 0,000 | 0,000 | 0,000 | 0,000 | 0,000 | 0,000 | 0,000 | 0,000 | 0,000 | 0,000 | 0,000 | 0,000 |
| Proteobacteria | *Pasteurella pneumotropica* | 0,000 | 0,000 | 0,000 | 0,000 | 0,094 | 0,094 | 0,000 | 0,000 | 0,000 | 0,000 | 0,000 | 0,000 | 0,000 | 0,000 |
| Proteobacteria | *Pelomonas saccharophila* | 0,000 | 0,000 | 0,000 | 0,000 | 0,192 | 0,192 | 0,000 | 0,000 | 0,000 | 0,000 | 0,000 | 0,000 | 0,000 | 0,000 |
| Proteobacteria | *Pseudomonas fluorescens* | 0,000 | 0,000 | 0,000 | 0,000 | 1,246 | 0,334 | 0,000 | 0,000 | 0,000 | 0,000 | 0,000 | 0,000 | 0,000 | 0,000 |
| Proteobacteria | *Pseudomonas sp,* | 0,107 | 0,107 | 0,000 | 0,000 | 0,660 | 0,152 | 0,000 | 0,000 | 0,000 | 0,000 | 0,000 | 0,000 | 0,000 | 0,000 |
| Proteobacteria | *Pseudomonas putida* | 0,000 | 0,000 | 0,000 | 0,000 | 0,174 | 0,122 | 0,000 | 0,000 | 0,000 | 0,000 | 0,000 | 0,000 | 0,000 | 0,000 |
| Proteobacteria | *Salmonella enterica* | 0,000 | 0,000 | 0,455 | 0,203 | 0,000 | 0,000 | 0,434 | 0,118 | 0,000 | 0,000 | 0,000 | 0,000 | 1,173 | 0,133 |
| Proteobacteria | *Shigella flexneri* | 0,000 | 0,000 | 1,338 | 0,543 | 0,000 | 0,000 | 1,217 | 0,375 | 0,000 | 0,000 | 0,160 | 0,095 | 2,317 | 0,310 |
| Proteobacteria | *Shigella sonnei* | 0,000 | 0,000 | 0,509 | 0,199 | 0,000 | 0,000 | 0,494 | 0,149 | 0,000 | 0,000 | 0,000 | 0,000 | 0,703 | 0,064 |
| Proteobacteria | *Shigella dysenteriae* | 0,000 | 0,000 | 0,055 | 0,055 | 0,000 | 0,000 | 0,056 | 0,056 | 0,000 | 0,000 | 0,000 | 0,000 | 0,113 | 0,113 |
| Proteobacteria | *Shigella boydii* | 0,000 | 0,000 | 0,120 | 0,080 | 0,000 | 0,000 | 0,186 | 0,124 | 0,000 | 0,000 | 0,000 | 0,000 | 0,000 | 0,000 |

| | | | | | | | | | | | | | | | |
|---|---|---|---|---|---|---|---|---|---|---|---|---|---|---|---|
| Proteobacteria | *Sphingomonas sp,* | 0,000 | 0,000 | 0,000 | 0,000 | 0,000 | 0,000 | 0,547 | 0,281 | 0,000 | 0,000 | 2,635 | 1,102 | 0,000 | 0,000 |
| Proteobacteria | *Sphingomonas ginsenosidimutans* | 0,000 | 0,000 | 0,000 | 0,000 | 0,000 | 0,000 | 0,069 | 0,069 | 0,000 | 0,000 | 0,348 | 0,202 | 0,000 | 0,000 |
| Proteobacteria | *Sphingomonas panni* | 0,000 | 0,000 | 0,000 | 0,000 | 0,000 | 0,000 | 0,000 | 0,000 | 0,000 | 0,000 | 0,000 | 0,000 | 0,000 | 0,000 |
| Proteobacteria | *Sphingomonas yunnanensis* | 0,000 | 0,000 | 0,000 | 0,000 | 0,000 | 0,000 | 0,000 | 0,000 | 0,000 | 0,000 | 0,365 | 0,365 | 0,000 | 0,000 |
| Proteobacteria | *Stenotrophomonas maltophilia* | 0,000 | 0,000 | 0,000 | 0,000 | 0,000 | 0,000 | 0,000 | 0,000 | 0,000 | 0,000 | 0,000 | 0,000 | 0,413 | 0,413 |
| Tenericutes | *Mycoplasma ovipneumoniae* | 1,927 | 0,910 | 2,071 | 0,810 | 2,480 | 0,825 | 1,074 | 0,553 | 3,555 | 2,506 | 3,055 | 1,398 | 0,000 | 0,000 |
| Tenericutes | *Mycoplasma hyopneumoniae* | 0,000 | 0,000 | 0,198 | 0,198 | 0,000 | 0,000 | 0,000 | 0,000 | 0,000 | 0,000 | 0,400 | 0,400 | 0,000 | 0,000 |
| Tenericutes | *Mycoplasma flocculare* | 0,000 | 0,000 | 0,131 | 0,131 | 0,000 | 0,000 | 0,000 | 0,000 | 0,000 | 0,000 | 0,000 | 0,000 | 0,000 | 0,000 |
| Verrucomicrobia | *Akkermansia muciniphila* | 0,000 | 0,000 | 1,194 | 0,794 | 0,164 | 0,164 | 1,436 | 0,713 | 0,000 | 0,000 | 1,400 | 1,400 | 0,450 | 0,450 |

**table S1. All species identified in stool samples from mice treated with TNBS alone or in combination with oxo- and allo-bile acid derivatives.**

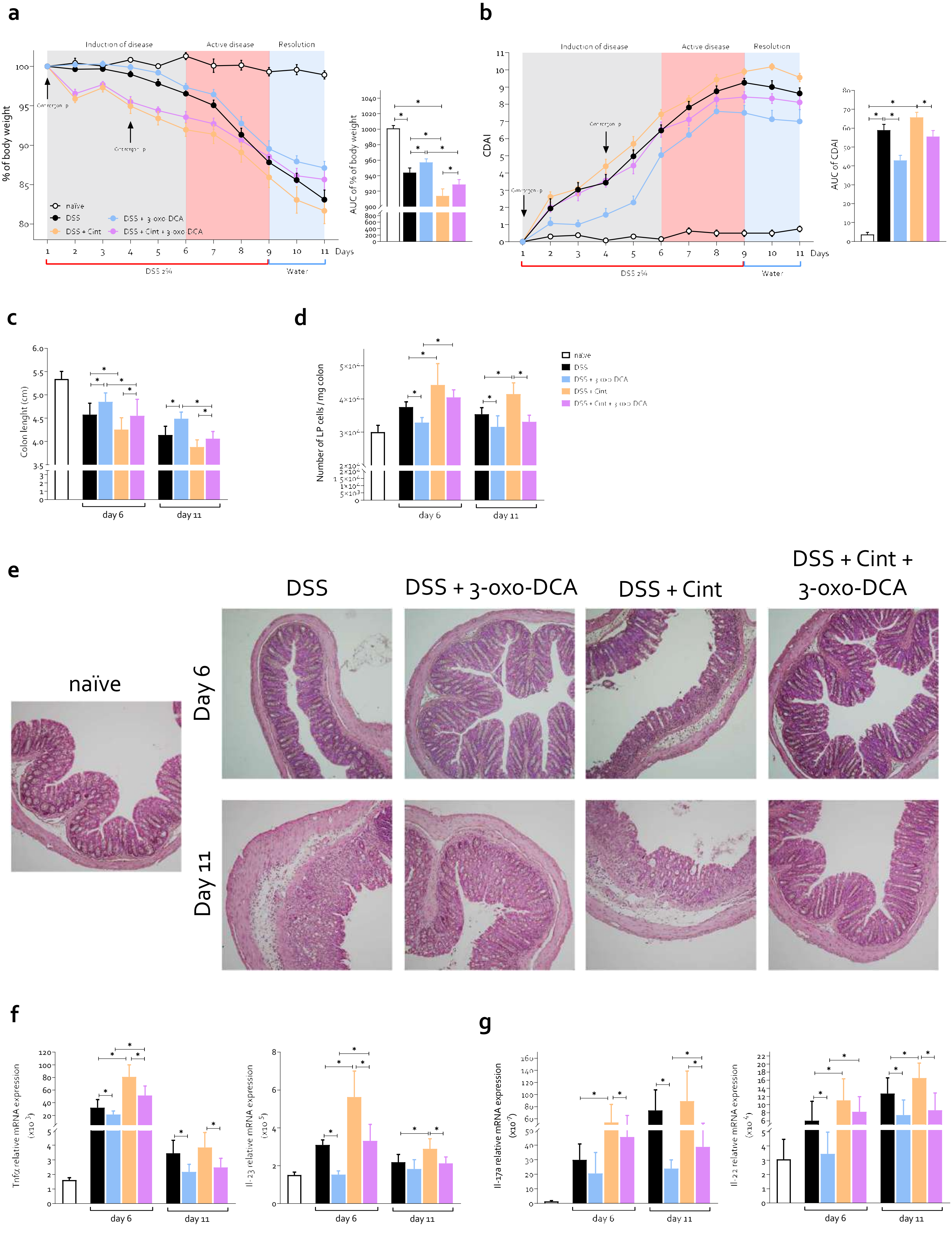
a
Induction of disease
Active disease
Resolution
% of body weight
naïve
DSS
DSS + Cint
DSS + 3-oxo-DCA
DSS + Cint + 3-oxo-DCA
Days
DSS 2%
Water
AUC of % of body weight
b
CDAI
AUC of CDAI
c
Colon lenght (cm)
day 6
day 11
d
Number of LP cells / mg colon
e
naïve
DSS
DSS + 3-oxo-DCA
DSS + Cint
DSS + Cint + 3-oxo-DCA
Day 6
Day 11
f
Tnfα relative mRNA expression (x10^-3)
Il-23 relative mRNA expression (x10^-5)
g
Il-17a relative mRNA expression (x10^-7)
Il-22 relative mRNA expression (x10^-4)

**Supplementary figure 5.** Colitis was induced by simultaneous administration of DSS + cintirorgon, a selective agonist of RORγt, in C57BL/6 wild-type mice. 2% DSS was administered in drinking water for 9 consecutive days. Cintirorgon was administered at a dose of 20 mg/kg via i.p. injection on day 1 and day 4. 3-oxo-DCA (10 mg/Kg/daily) was administered by gavage (o.s.) from day 1 to the end of the experiments. The mice were sacrificed at two time points: at the end of the disease induction phase (day 6) or during the resolution phase on day 11 (DSS administration was suspended on day 9). The severity of the disease at day 6 and 11 was assessed by: (**a**) percentage of body weight change and area under the curve (AUC) of body weight trends; (**b**) Colitis Disease Activity Index (CDAI) and AUC of CDAI, (**c**) measurement of colon length and (**d**) ratio between *lamina propria* cells and colon weight (mg); (**e**) microscopic analysis of the colon using hematoxylin and eosin staining (magnification 10×).(**f**-**g**) Relative mRNA expression of (**f**) Tnfα and Il-23 and (**g**) IL-17a and IL-22 measured in the *lamina propria* cells of the colon. Data are normalized to GAPDH mRNA. Data shown are the mean ± SEM of 4-12 mice per group. Statistical significance was assessed by 1-way ANOVA $*p \leq 0.05$.

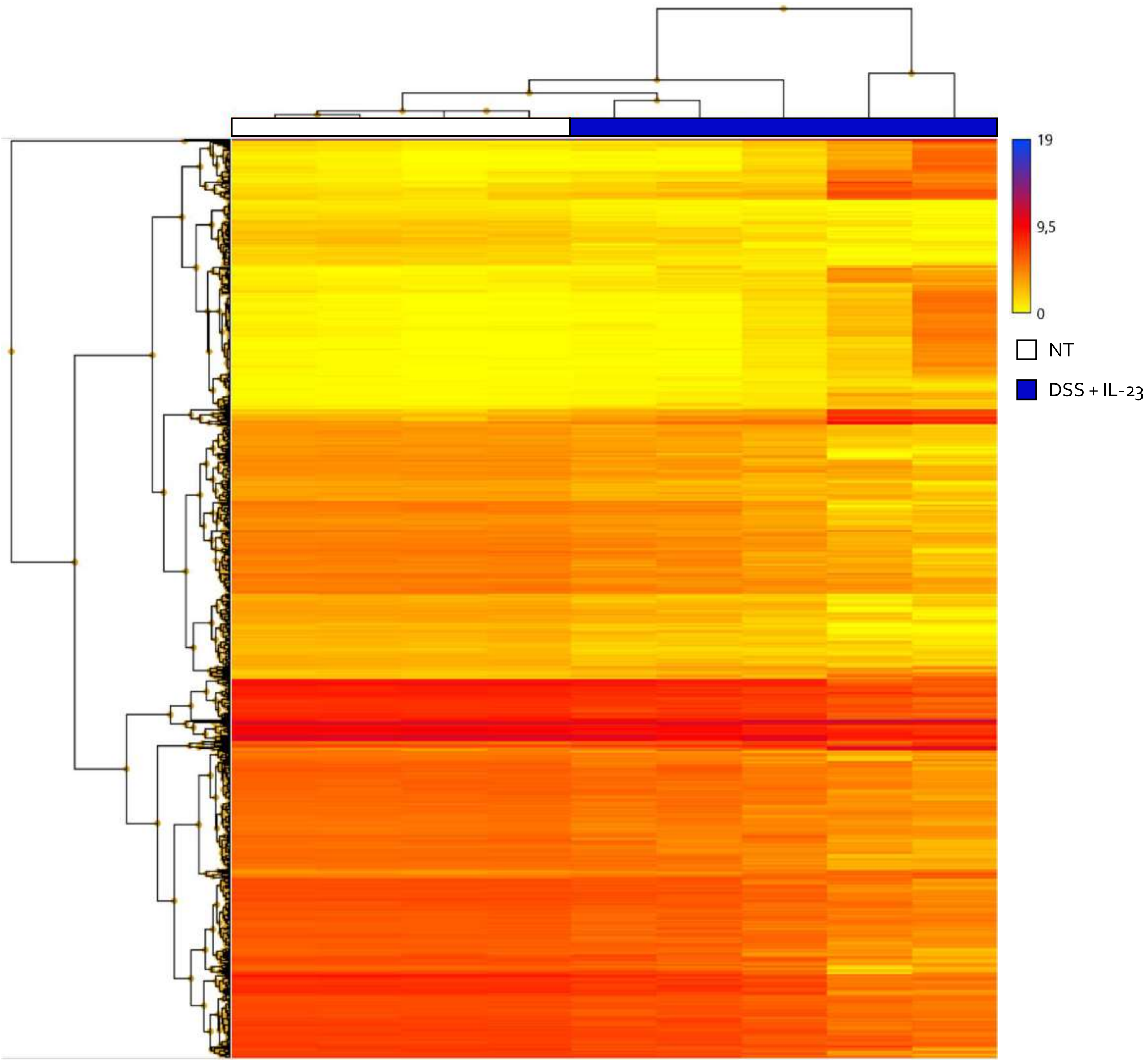


**Supplementary figure 6.** Colitis was induced by simultaneous administration of DSS + IL-23 in C57BL/6 wild-type mice. 2% DSS was administered in drinking water for 6 consecutive days. IL-23 was administered at a dose of 500 ng/mouse via i.p. injection daily. 3-oxo-DCA (10 mg/Kg/daily) was administered by o.s. from day 1 to the end of the experiments. The mice were sacrificed at day 6. Hierarchically clustered heatmap of differentially expressed genes between Not treated samples and DSS/IL23 treated samples. The transition from yellow to blue strips represents an increase in gene expression levels**.** Abbreviations: NT, not treated; DSS, Dextran sulfate sodium; IL23, interleukin-23.

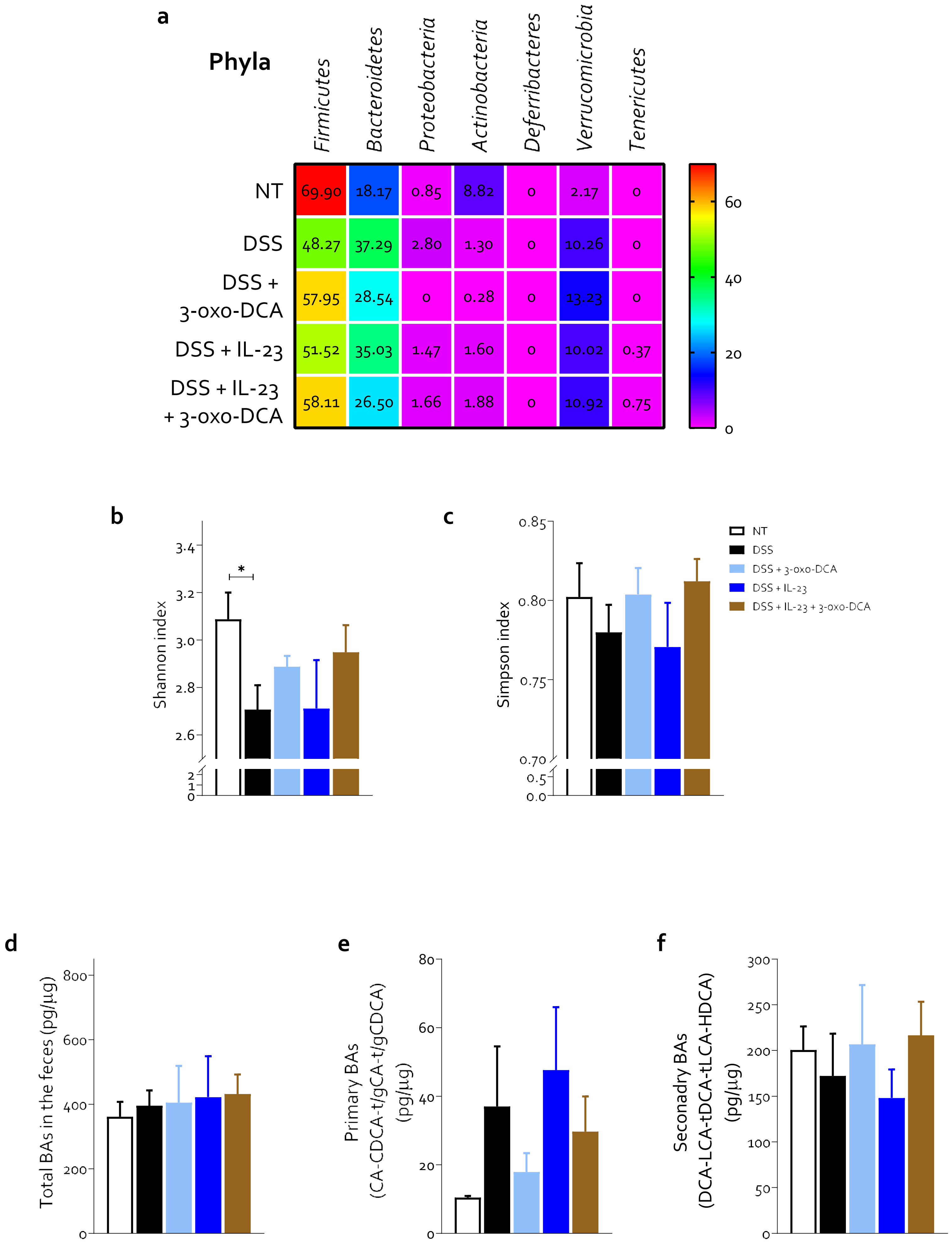
a
Phyla
Firmicutes
Bacteroidetes
Proteobacteria
Actinobacteria
Deferribacteres
Verrucomicrobia
Tenericutes
NT
69.90
18.17
0.85
8.82
0
2.17
0
DSS
48.27
37.29
2.80
1.30
0
10.26
0
DSS + 3-oxo-DCA
57.95
28.54
0
0.28
0
13.23
0
DSS + IL-23
51.52
35.03
1.47
1.60
0
10.02
0.37
DSS + IL-23 + 3-oxo-DCA
58.11
26.50
1.66
1.88
0
10.92
0.75
60
40
20
0
b
Shannon index
*
3.4
3.2
3.0
2.8
2.6
2
1
0
c
Simpson index
0.85
0.80
0.75
0.70
0.5
0.0
NT
DSS
DSS + 3-oxo-DCA
DSS + IL-23
DSS + IL-23 + 3-oxo-DCA
d
Total BAs in the feces (pg/µg)
800
600
400
200
0
e
Primary BAs
(CA-CDCA-t/gCA-t/gCDCA)
(pg/µg)
80
60
40
20
0
f
Secondary BAs
(DCA-LCA-tDCA-tLCA-HDCA)
(pg/µg)
300
250
200
150
100
50
0

**Supplementary figure 7.** Colitis was induced by simultaneous administration of DSS + IL-23 in C57BL/6 wild-type mice. 2% DSS was administered in drinking water for 6 consecutive days. IL-23 was administered at a dose of 500 ng/mouse via i.p. injection daily. 3-oxo-DCA (10 mg/Kg/daily) was administered by o.s. from day 1 to the end of the experiments. The mice were sacrificed at day 6. Data shown are: (**a**) relative abundance of different Phyla expressed as percent of mapped reads in each experimental group and measurement of community heterogeneity using (**b**) Shannon and (**c**) Simpson indexes at family level evaluated in fecal samples. Fecal levels of (**d**) total bile acids, (**e**) primary bile acids (CA-CDCA-t/gCA-t/gCDCA) and (**f**) secondary bile acids (DCA-LCA-tDCA-tLCA-HDCA). Data shown are the mean ± SEM of 3-7 mice per group. Statistical significance was assessed by 1-way ANOVA $*p \leq 0.05$. Abbreviations: NT, not treated DSS, Dextran sulfate sodium; IL23, interleukin-23; DCA, deoxycholic acid; t, tauro; g, glyco; CA, cholic acid; CDCA, chenodeoxycholic acid; LCA, lithocholic acid; HDCA, hyodeoxycholic acid.

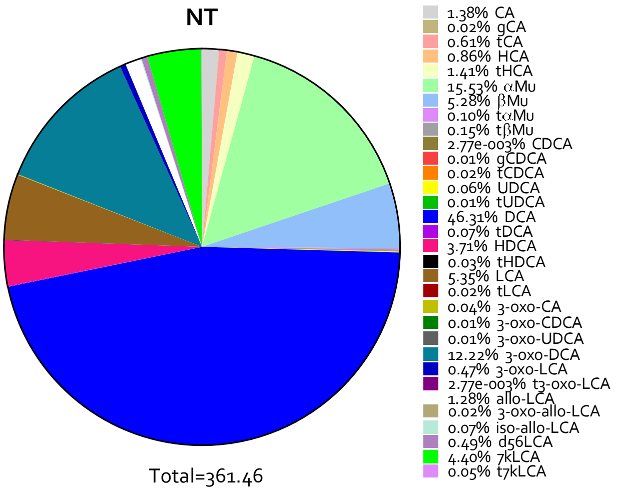
NT
1.38% CA
0.02% gCA
0.61% tCA
0.86% HCA
1.41% tHCA
15.53% αMu
5.28% βMu
0.10% tαMu
0.15% tβMu
2.77e-003% CDCA
0.01% gCDCA
0.02% tCDCA
0.06% UDCA
0.01% tUDCA
46.31% DCA
0.07% tDCA
3.71% HDCA
0.03% tHDCA
5.35% LCA
0.02% tLCA
0.04% 3-oxo-CA
0.01% 3-oxo-CDCA
0.01% 3-oxo-UDCA
12.22% 3-oxo-DCA
0.47% 3-oxo-LCA
2.77e-003% t3-oxo-LCA
1.28% allo-LCA
0.02% 3-oxo-allo-LCA
0.07% iso-allo-LCA
0.49% d56LCA
4.40% 7kLCA
0.05% t7kLCA
Total=361.46

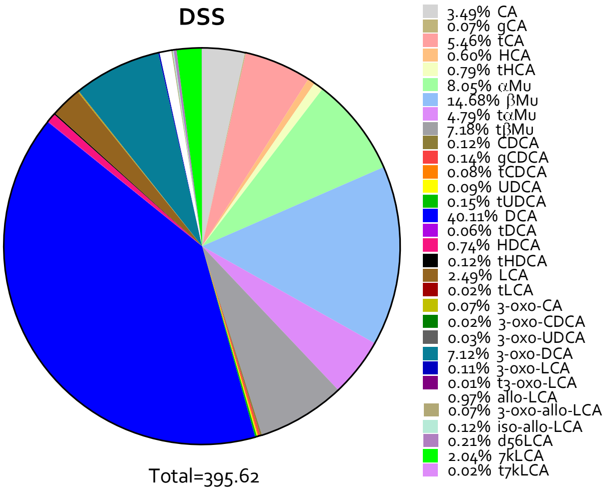
DSS
3.49% CA
0.07% gCA
5.46% tCA
0.60% HCA
0.79% tHCA
8.05% αMu
14.68% βMu
4.79% tαMu
7.18% tβMu
0.12% CDCA
0.14% gCDCA
0.08% tCDCA
0.09% UDCA
0.15% tUDCA
40.11% DCA
0.06% tDCA
0.74% HDCA
0.12% tHDCA
2.49% LCA
0.02% tLCA
0.07% 3-oxo-CA
0.02% 3-oxo-CDCA
0.03% 3-oxo-UDCA
7.12% 3-oxo-DCA
0.11% 3-oxo-LCA
0.01% t3-oxo-LCA
0.97% allo-LCA
0.07% 3-oxo-allo-LCA
0.12% iso-allo-LCA
0.21% d56LCA
2.04% 7kLCA
0.02% t7kLCA
Total=395.62

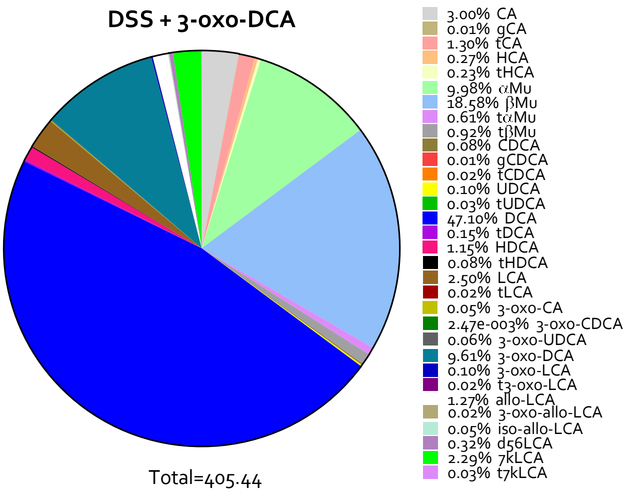
DSS + 3-oxo-DCA
3.00% CA
0.01% gCA
1.30% tCA
0.27% HCA
0.23% tHCA
9.98% αMu
18.58% βMu
0.61% tαMu
0.92% tβMu
0.08% CDCA
0.01% gCDCA
0.02% tCDCA
0.10% UDCA
0.03% tUDCA
47.10% DCA
0.15% tDCA
1.15% HDCA
0.08% tHDCA
2.50% LCA
0.02% tLCA
0.05% 3-oxo-CA
2.47e-003% 3-oxo-CDCA
0.06% 3-oxo-UDCA
9.61% 3-oxo-DCA
0.10% 3-oxo-LCA
0.02% t3-oxo-LCA
1.27% allo-LCA
0.02% 3-oxo-allo-LCA
0.05% iso-allo-LCA
0.32% d56LCA
2.29% 7kLCA
0.03% t7kLCA
Total=405.44

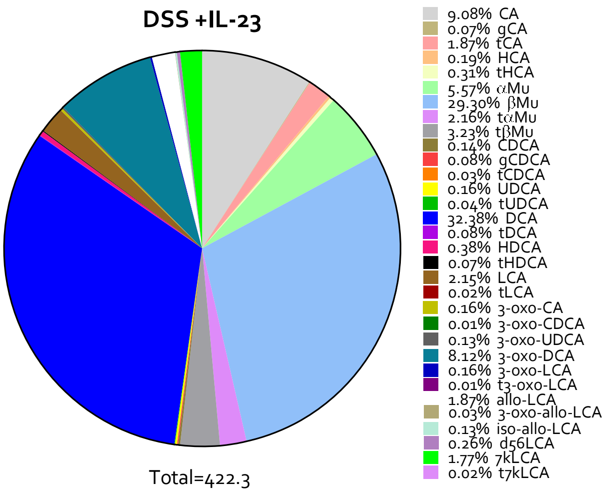
DSS +IL-23
9.08% CA
0.07% gCA
1.87% tCA
0.19% HCA
0.31% tHCA
5.57% αMu
29.30% βMu
2.16% tαMu
3.23% tβMu
0.14% CDCA
0.08% gCDCA
0.03% tCDCA
0.16% UDCA
0.04% tUDCA
32.38% DCA
0.08% tDCA
0.38% HDCA
0.07% tHDCA
2.15% LCA
0.02% tLCA
0.16% 3-oxo-CA
0.01% 3-oxo-CDCA
0.13% 3-oxo-UDCA
8.12% 3-oxo-DCA
0.16% 3-oxo-LCA
0.01% t3-oxo-LCA
1.87% allo-LCA
0.03% 3-oxo-allo-LCA
0.13% iso-allo-LCA
0.26% d56LCA
1.77% 7kLCA
0.02% t7kLCA
Total=422.3

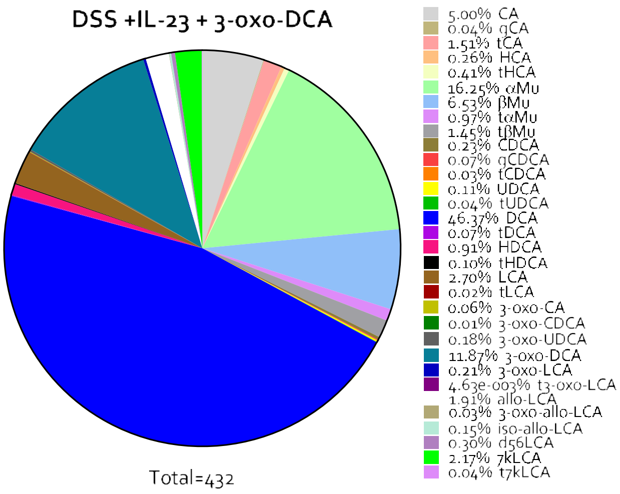
DSS +IL-23 + 3-oxo-DCA
5.00% CA
0.04% gCA
1.51% tCA
0.26% HCA
0.41% tHCA
16.25% αMu
6.53% βMu
0.97% tαMu
1.45% tβMu
0.23% CDCA
0.07% gCDCA
0.03% tCDCA
0.11% UDCA
0.04% tUDCA
46.37% DCA
0.07% tDCA
0.91% HDCA
0.10% tHDCA
2.70% LCA
0.02% tLCA
0.06% 3-oxo-CA
0.01% 3-oxo-CDCA
0.18% 3-oxo-UDCA
11.87% 3-oxo-DCA
0.21% 3-oxo-LCA
4.63e-003% t3-oxo-LCA
1.91% allo-LCA
0.03% 3-oxo-allo-LCA
0.15% iso-allo-LCA
0.30% d56LCA
2.17% 7kLCA
0.04% t7kLCA
Total=432

**Supplementary figure 8.** Colitis was induced by simultaneous administration of DSS + IL-23 in C57BL/6 wild-type mice. 2% DSS was administered in drinking water for 6 consecutive days. IL-23 was administered at a dose of 500 ng/mouse via i.p. injection daily. 3-oxo-DCA (10 mg/Kg/daily) was administered by o.s. from day 1 to the end of the experiments. The mice were sacrificed at day 6. Histogram of fecal content of total bile acids evaluated in each experimental group. Abbreviations: NT, not treated; DSS, Dextran sulfate sodium; T, tauro; G, glyco; LCA, lithocholic acid; DCA, deoxycholic acid; CDCA, chenodeoxycholic acid; HDCA, hyodeoxycholic acid; UDCA, ursodeoxycholic acid; HCA, hyocholic acid; αMu, alpha-muricholic acid; βMu, beta-muricholic acid; 7k, 7keto; CA, cholic acid.
